# Unraveling the pathway of Copper Delivery to Cytochrome *c* oxidases in the Free-Living Bacterium *Caulobacter vibrioides*

**DOI:** 10.1101/2025.09.05.674415

**Authors:** Hala Kasmo, Jacquie Abolia Tepusa, Rubén Garcia-Dominguez, Chloe Piette, Marc Dieu, Damien Devos, Jean-Yves Matroule

## Abstract

Copper (Cu) is an essential micronutrient that serves as a cofactor for many enzymes but becomes toxic when present in excess. In most bacteria, CopA-like P1B-type ATPases mediate Cu detoxification by exporting cytoplasmic Cu to the periplasm or extracellular environment.

In this study, we show that *Caulobacter vibrioides* lacks a canonical CopA-like ATPase but encodes a single FixI/CcoI-type Cu-transporting ATPase, previously implicated in Cu delivery to the *cbb*□-type cytochrome *c* oxidase (Cox) in species such as *Rhodobacter capsulatus*. *C. vibrioides* harbors two terminal cytochrome *c* oxidases in its cytoplasmic membrane: an *aa*□-type and a *cbb*□-type Cox. We also demonstrate that the activity of *cbb*□-Cox requires the FixI-type Cu transporter and the periplasmic Cu chaperone PccA. In contrast, *aa*□-Cox activity depends on PccA and the inner membrane-bound protein CtaG. Since the mechanism of Cu acquisition for *aa*□-Cox remains largely unknown, we conducted a genetic screen and identified a novel outer membrane TonB-dependent receptor (TccA) that is specifically required for *aa*□-Cox function.

We also showed that *cbb*□-Cox is upregulated under microaerobic conditions, possibly such as those encountered on solid media where O_2_ diffusion is limited. Under normoxic conditions, the expression and the activity of *cbb*□-Cox decrease, and *aa*□-Cox becomes the predominant terminal oxidase. These findings demonstrate that *C. vibrioides* differentially utilizes its Cox enzymes in response to O_2_ availability and relies on a distinct Cu trafficking pathway for their maturation, including an outer membrane component that has not been previously described in bacterial Cu homeostasis.

## Introduction

Copper (Cu) is a redox-active metal playing a key role in several biological processes, including respiration and defense against oxidative stress. Its unique ability to switch between the reduced Cu(I) and oxidized Cu(II) ionic states makes it a vital catalytic cofactor in redox enzymes such as cytochrome oxidases (Cox) and Cu-Zn superoxide dismutases (1, 2). However, the same redox properties render Cu toxic at elevated concentrations, requiring tightly regulated systems for its uptake, trafficking, and efflux (1, 2).

The primary defense against cytoplasmic Cu overload in most bacteria is the P1B1-type ATPase CopA, a membrane-bound transporter that actively exports Cu(I) from the cytoplasm to the periplasm, where it can be chelated, utilized, or further expelled (3, 4). The activity of CopA often relies on the small Cu chaperone CopZ(5, 6), which binds Cu(I) in the cytoplasm through conserved cysteine motifs (CxxC) and delivers it to CopA (7), thereby buffering intracellular Cu and facilitating its specific export from the cell. However, not all P1B1-type ATPases ensure Cu detoxification. In several bacterial species, including *Rhodobacter capsulatus* and *Pseudomonas aeruginosa* (8, 9), dedicated FixI/CcoI-type P1B-type ATPases mediate Cu delivery specifically for the biogenesis of the terminal *cbb_3_*-Cox enzymes in the respiratory chain (10).

Terminal oxidases of the heme–Cu oxidase (HCO) superfamily, including *aa*□*-*type and *cbb*□*-*type Cox, catalyze the four-electron reduction of O_2_ into H_2_O, a critical step in aerobic respiration (11). These enzymes require precise Cu insertion into their catalytic centers for activity. The *aa*□*-*type Cox has a low affinity for O_2_ and typically contains three core subunits: CoxA (subunit I), CoxB (subunit II), and CoxC (subunit III)(11–13). CoxA harbors the binuclear catalytic center composed of heme *a*, heme *a*□-CuB site, while CoxB harbors the binuclear Cu center CuA. It acts as the primary electron acceptor from cytochrome *c*, transferring electrons to the heme *a* and then to the heme *a_3_*-CuB binuclear center, where O□ is reduced to H_2_O. The proper assembly of the CuA and CuB sites is essential for the Cox enzyme function and requires the coordinated action of multiple accessory proteins (11, 14, 15). In bacteria such as *Rhodobacter sphaeroides* (16), the maturation of the *aa*□- type Cox relies on the Sco-like and PCu(A)C chaperones for Cu delivery to CuA and CuB centers, while CtaG is required for CuB center. In *Paracoccus denitrificans*, Cu delivery to the *aa_3_*-type Cox involves multiple chaperone proteins (17). Besides CtaG, which is implicated in the insertion of Cu into the CuB center, two Sco-like proteins have been identified. ScoB plays a critical role in Cu delivery to the CuA center, whereas ScoA and the two PCuAC-like proteins appear to have less direct effects on the enzyme activity (17). The *cbb*□-type Cox is a high-affinity terminal oxidase composed of at least three subunits: CcoN, CcoO, and CcoP. The catalytic subunit, CcoN, contains heme *b* and heme *b*□*–*CuB center. The subunits CcoP and CcoO serve as periplasmic electron acceptors that transfer electrons from cytochrome *c* to the catalytic subunit CcoN, where O_2_ is reduced to H_2_O. Cu plays a key role in the formation of the CuB site, the assembly of which requires specific trafficking pathways. In *R. capsulatus*, the inner membrane (IM) transporter CcoA, a member of the major facilitator superfamily (MFS), mediates Cu uptake into the cytoplasm (18). Once inside, CcoI exports Cu into the periplasm, where the Cu chaperones PccA and SenC coordinate the final steps of Cu delivery to the CuB center (8). PccA binds Cu in the periplasm and likely transfers it to SenC, which directly participates in the metallation of the CuB center (19). The coordinated action of these components is essential for *cbb*□-Cox activity, and mutations in any of the associated genes result in severe defects in enzyme assembly and respiratory function (10, 19).

In *Bradyrhizobium japonicum*, the maturation of *aa*□-Cox relies on CtaG-like protein CoxG and ScoI, specifically required for the assembly of the CuB center in subunit I and the CuA center in subunit II, respectively (20). Deletion of either *coxG* or *scoI* lead to a complete loss of *aa*□-Cox assembly and activity in the cytoplasmic membrane, while the *cbb*□-Cox remains unaffected. This highlights the distinct Cu-handling pathways that exist for these two types of terminal oxidases (20). In addition, the PcuAC-like protein PcuC plays a crucial role in the biogenesis of both *aa*□- and *cbb*□-Cox, especially under Cu-limiting conditions (21). Studies in *R. sphaeroides*, *B. japonicum,* and other α-proteobacteria have significantly advanced our understanding of Cox assembly and electron transfer, particularly regarding the structural organization of the *aa_3_*- and *cbb_3_*-type Coxs and the roles of the CuA and CuB metal centers. However, despite these insights, Cu transport, delivery pathways, and Cox biogenesis in other α-proteobacteria remain largely uncharacterized. *C. vibrioides* is an important model for studying bacterial cell cycle regulation, environmental adaptation, and metal homeostasis (22). As an oligotrophic, strictly aerobic, free-living alphaproteobacterium, it is constantly exposed to fluctuating O_2_ and metal levels, making it ideal for investigating how bacteria coordinate respiration and metal transport. *C. vibrioides* encodes four terminal oxidases: Cyt*bd* and Cyt*bo*□, *aa*□-Cox, and *cbb*□-Cox, enabling it to respire under a wide range of O_2_ concentrations. Despite the widespread distribution of *aa*□-type Cox in aerobic bacteria, whether a specific Cu transporter exists for this enzyme class remains unclear. No dedicated Cu import system has yet been identified for *aa*□-Cox, and the mechanisms of Cu acquisition remain poorly understood. This study addressed this gap by identifying TccA, a TonB-dependent outer membrane transporter, as crucial for full *aa*□-Cox activity, likely by mediating Cu uptake into the periplasm and delivering it to the CuA site, revealing a novel role for such receptors in bacterial Cu acquisition. In contrast, *cbb*□-Cox assembles via an alternative Cu pathway under microoxic conditions, highlighting O_2_-dependent respiratory flexibility in *C. vibrioides* and suggesting broader diversity in bacterial Cu utilization mechanisms.

## Results

### *Caulobacter vibrioides* P1B1-type ATPase is not required for Cu detoxification

Using the Basic Local Alignment Search Tool (BLAST) to find homologous sequences, we queried the *C.*□*vibrioides* NA1000 proteome (23) with the *E.*□*coli* CopA protein sequence and identified a single P1B1-type ATPase, hereafter referred to as FixI . The multiple sequence alignments of FixI with various well-studied P1B1-type ATPases highlighted two conserved Cu-binding sites of CopA (Site I: C398, C400, Y681, and Site II: N682, M703, S706) (Fig. 1A). The whole sequence alignment is indicated in Fig. S1. The Δ*fixI* mutant displayed a WT-like growth pattern in liquid-rich (PYE) medium, indicating that FixI is not required for *C. vibrioides* survival under control condition (Fig. 1B, upper panel). No increased sensitivity was observed when the Δ*fixI* mutant was exposed to moderate Cu stress (150 µM) in liquid and solid medium (Fig. 1B, lower panel and Fig. 1C). Consistent with these data, no significant difference in the intracellular Cu ions content could be measured between the WT and the Δ*fixI* strains by atomic absorption spectroscopy (AAS) (Fig. 1D). Similarly, the Δ*fixI* mutant did not show any increased sensitivity to Cobalt (Co), Nickel (Ni), Zinc (Zn) and Cadmium (Cd) in liquid and solid medium (Fig. S2A and Fig. S2B).

**Figure 1.**
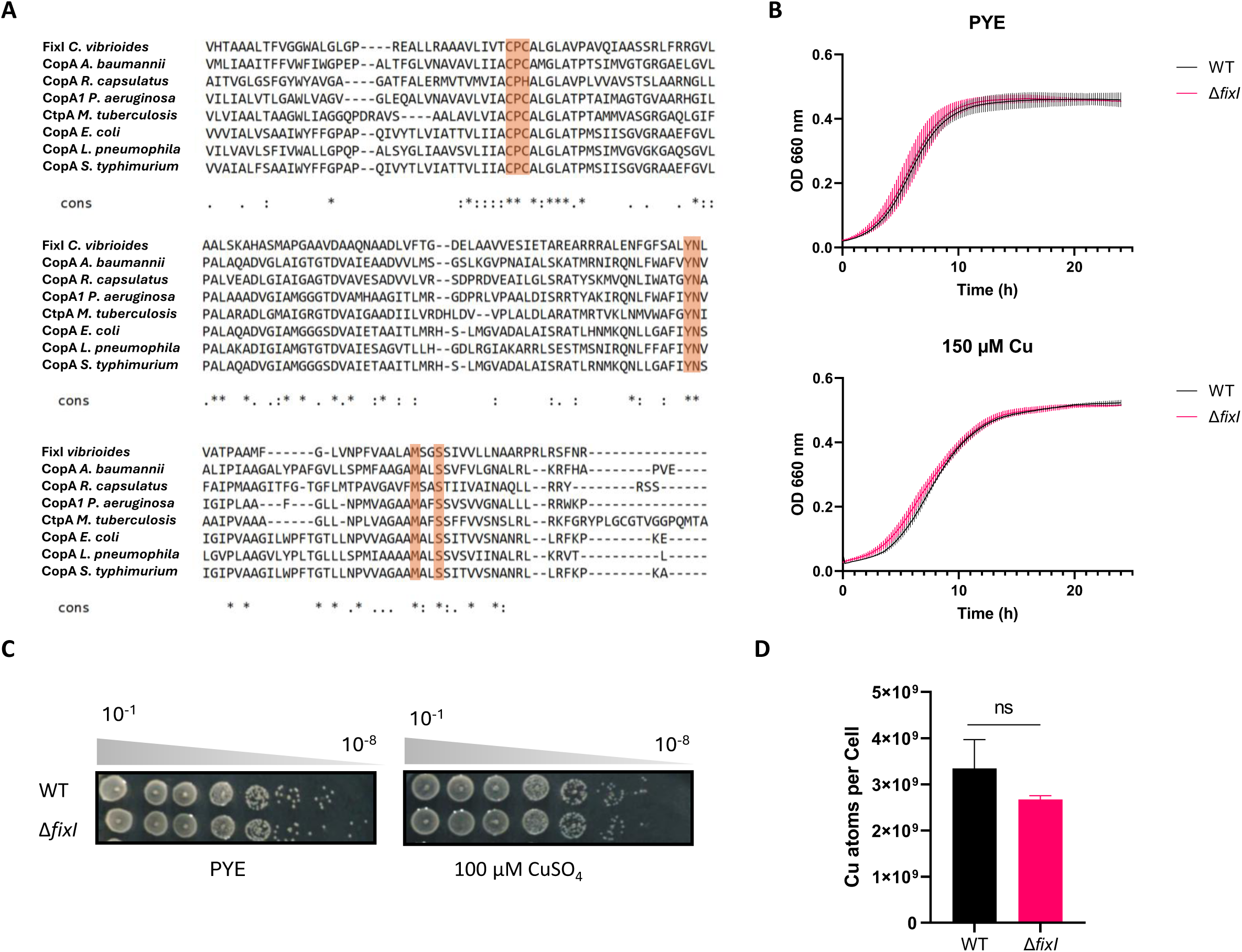
The single P1B1-type ATPase of *Caulobacter vibrioides* is not required for Cu detoxification. **A**. Amino acid sequence alignment of FixI from *C. vibrioides* with its homologs CopA from *Acinetobacter baumannii*, CopA from *Rhodobacter capsulatus*, FixI *Pseudomonas aeruginosa*, CtpA *Mycobacterium tuberculosis,* CopA *Escherichia coli*, CopA *Legionella pneumophila,* and CopA *Salmonella typhimurium*. The conserved Cu-binding sites, including the CPC motif, are highlighted by orange boxes. **B**. Growth profiles of WT and Δ*fixI* strains at OD_660_ in control (top) and moderate Cu stress (bottom) conditions. Mean ± SD, at least three biological replicates. **C**. Viability assay on rich (PYE) in control and Cu excess conditions with 100 μM CuSO_4_, respectively. Plates were incubated for 48 h at 30 °C before imaging. Biological replicates = 3. **D**. Number of Cu atoms per cell in the control condition and after 30 min of 175 µM CuSO_4_ exposure. Mean ± SD, at least three biological replicates. P-values were calculated using a t-test (\**p* < 0.05, ** *p* < 0.01, *** *p* < 0.001 and \*\*\*\**p* < 0.0001) **(Table S4)**..

The lack of Cu sensitivity of the Δ*fixI* mutant may result from a functional redundancy with other Cu transporters that could compensate for the loss of FixI. The analysis of the *C. vibrioides* genome highlighted a P1B4-type ATPase, referred to as ZctP in this study. P1B4-type ATPases are best known for their role in the translocation of a wide range of substrates (24). To test whether ZctP plays a role in Cu efflux in *C. vibrioides*, a single Δ*zctP* mutant and a double Δ*fixI*Δ*zctP* mutant were generated, and their growth was monitored under moderate Cu stress in both liquid and solid media. Both Δ*zctP* and Δ*fixI*Δ*zctP* mutants displayed WT-like growth profiles under Cu stress (Fig. S2C and Fig. S2D) but not under Zn stress (additional information 1).

Together, these data suggest that *C. vibrioides* lacks the P1B-type ATPase required for Cu detoxification.

### FixI is required for *cbb***□**-Cox full activity

FixI is encoded by an operon involved in the assembly and activity of *cbb*□-Cox (Fig. 2A, upper panel). Given that the proteins encoded by genes within the same operon often participate in related biological processes, we hypothesized that FixI is required for Cu ion delivery to the CuB catalytic center of CcoN. The Cox activity of the WT, Δ*fixI,* and Δ*cbb*□ strains was monitored on plate by using the Cox-specific NADI staining (25). The WT strain was NADI^+^ (stained dark blue within 30 seconds), while both ΔfixI and Δ*cbb*□ mutants exhibited a NADI^slow^ phenotype (stained dark blue in 30 minutes), suggesting that Cox activity relies on FixI (Fig. 2B). The introduction of an exogenous copy of FixI under the constitutive p*lac* promoter on the pMR10 vector in the Δ*fixI* mutant (Δ*fixI*+p*fixI*) restored the NADI^+^ phenotype, attesting the correct Cox activity and the absence of polar effects.

**Figure 2.**
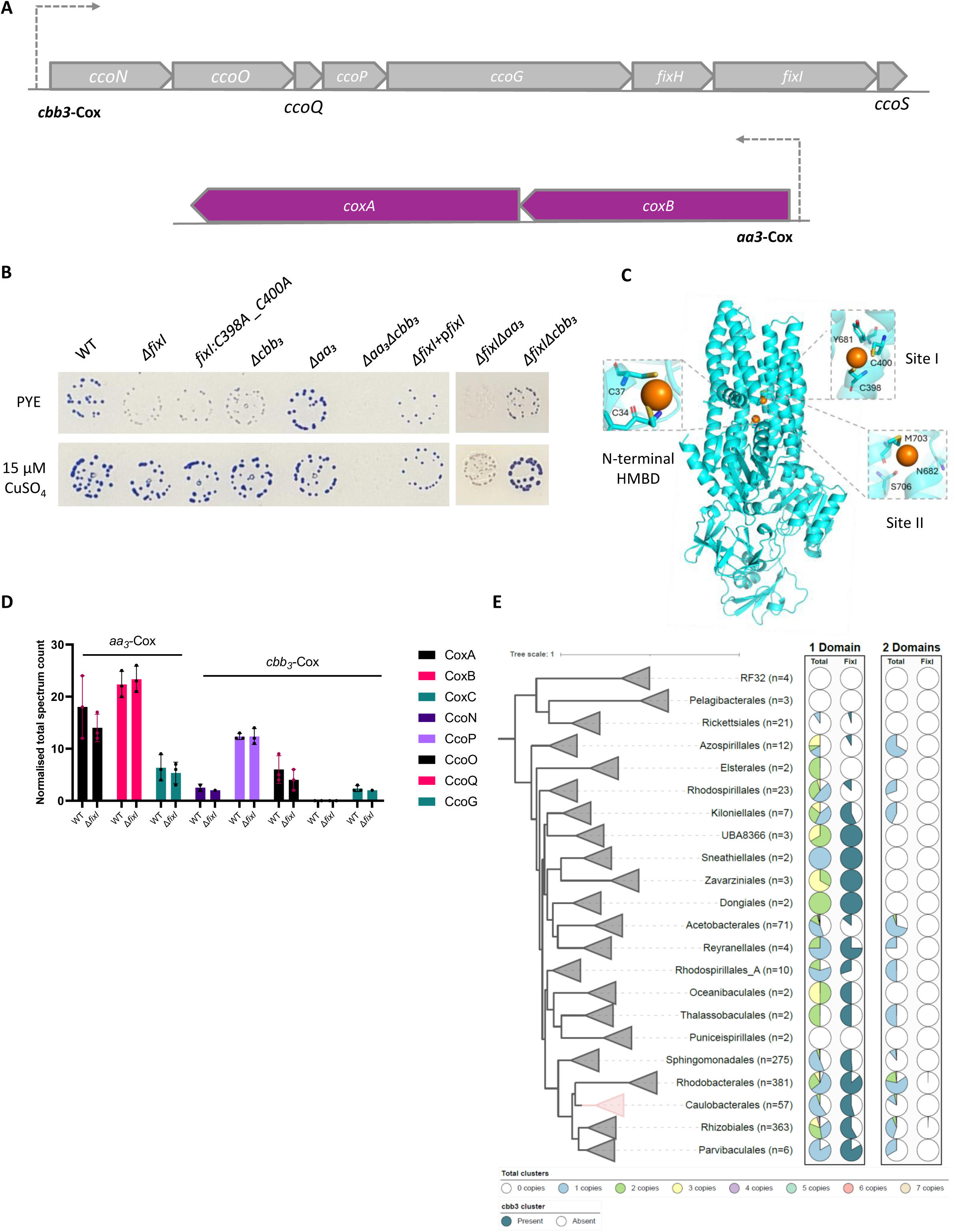
The full activity of *cbb*□-Cox is dependent on the Cu delivery through FixI. **A.** Genomic organization of the *cbb*□-Cox and *aa*□-Cox operon (Top) in *Caulobacter vibrioides* NA1000. **B.** Growth and NADI phenotypes of colonies of *C. vibrioides* WT strain and Δ*fixI*, *fixI::C398A_C400A*, Δ*cbb*□, Δ*aa*□, and Δ*aa*□Δ*cbb*□ mutants, together with those of Δ*fixI*Δ*cbb*□and Δ*fixI*Δ*aa*□mutants. Cells were grown aerobically at 30°C on a PYE medium and PYE supplemented with 15 µM CuSO_4_, and the presence of Cox activity was visualized by NADI staining. Colonies that contain wild-type levels of Cox activity turn dark blue within a few seconds (NADI^+^), while those that have low or no Cox activity show lighter blue (NADI^slow^) or no blue staining (NADI^-^) upon longer exposure, respectively. Biological replicates = 3 (entire plates in Fig. S11) **C**. Prediction by AlphaFold 3 of FixI 3D structure. The model was visualized using Pymol. **D.** Normalized total spectrum count of peptides of *aa*□-Cox subunits and *cbb*□-Cox subunits in both WT and Δ*fixI* strains grown in PYE medium, measured by LC-MS. Individual values and means are represented. Biological replicates = 3. *P*-values were calculated using a t-test. (\**p* < 0.05, ** *p* < 0.01, *** *p* < 0.001 and \*\*\*\**p* < 0.0001) **(Table S4)**. **E.** A collapsed subtree of the GTDB bacterial phylogeny at the order level, restricted to Alphaproteobacteria (n = 1,255 genomes with Pfam annotations). Two P-type ATPase domain architectures were identified: a 1-domain variant (PF00403–PF00122–PF00702) and a 2-domain variant (PF00403–PF00403–PF00122–PF00702), differing by a duplication of the N-terminal heavy metal-associated domain (PF00403). For each variant, the first column shows the proportion of genomes in each order carrying 0 to 7 copies (color-coded by copy number), and the second column shows the proportion of genomes encoding a *cbb*□-type cytochrome *c* oxidase gene cluster, potentially linked to FixI function.

The structure prediction of FixI by AlphaFold 3 (26) highlighted three potential Cu-binding sites located within the transmembrane domain (Site I and Site II) and at the N-terminal of the protein (Site III; C34xxC37) known as Heavy-metal binding domain (HMBD) (Fig. 2C) (27). In *E. coli*, the cysteine residues of the CPC motif (Site I in *C. vibrioides*) are required for CopA function in Cu resistance (3). The substitution of the corresponding cysteines (C398 and C400) into alanines in FixI phenocopied the NADI^slow^ phenotype of the Δ*fixI* mutant, confirming their key role in FixI activity (Fig. 2B). Interestingly, supplementing the medium with 15 µM CuSO_4_ restored the full Cox activity in the Δ*fixI*, *fixI*: C398A_C400A and Δ*cbb*□ mutants (Fig. 2B).

The NADI^slow^ phenotype observed in the aforementioned mutants may be related to the activity of the *aa*□-Cox (Fig. 2A). Consistent with this hypothesis, no Cox activity (NADI□) was detected in the double Δ*aa*□Δ*cbb*□ mutant (Fig. 2B). However, unlike the Δ*cbb*□ mutant, the Δ*aa*□ mutant exhibited a NADI□ phenotype, suggesting that *cbb*□-Cox is the dominant Cox in solid PYE medium (Fig. 2B). One could imagine that the bypass of the NADI^slow^ phenotype of Δ*cbb*□ by extracellular CuSO_4_ results from a boosted activity of *aa*□-Cox. Accordingly, the Δ*aa*□Δ*cbb*□ mutant remained NADI□ when excess CuSO_4_ in the medium (Fig. 2B). To verify whether the residual activity in the NADI^slow^ phenotype is due to the presence of steady Cu ions in the control medium, Cu was chelated using the specific Cu chelator bathocuproine sulfonate (BCS) in the minimal M2G medium. The NADI stain in M2G showed very faint activity; however, when 600 µM BCS was added, all tested strains, including the WT, were NADI^-^, indicating that both *aa_3_* and *cbb_3_*-Cox are inactive due to the lack of Cu ions in the media (Fig. S5A).

*C. vibrioides* is a strict aerobic bacterium, raising the question of how it remains viable in the absence of both *cbb*□- and *aa*□-Cox enzymes. The genome analysis *C. vibrioides* revealed the presence of the cytochrome *bd*-type quinol oxidase (Cyt*bd*) and cytochrome *bo3* quinol oxidase (Cyt*bo3*) respiratory terminal oxidases (Fig. S5B). The abundance of these proteins was not altered in the absence of both *aa*□ and *cbb*□-Cox compared to the WT strain (Fig. S5C).

To determine whether FixI delivers Cu ions to both *cbb*□- and *aa*□-Cox enzymes, double Δ*fix*Δ*cbb*□ and Δ*fixI,aa*□ mutants were generated, and their respective Cox activity was measured. The Δ*fixI*Δ*cbb*□ mutant displayed a NADI^slow^ phenotype, similar to the single Δ*fixI* and Δ*cbb*□ mutants (Fig. 2B). In contrast, the Δ*fixI*Δ*aa*□ mutant exhibited a NADI□ phenotype, indicating a complete loss of Cox activity despite the presence of an intact *cbb*□*-*Cox (Fig. 2B). These findings suggest that FixI is required for *cbb*□-Cox activity but not for *aa*□-Cox activity. Notably, supplementation with extracellular CuSO_4_ failed to restore *cbb*□-Cox activity in the absence of FixI, supporting the idea that the delivery of Cu ions to the *cbb*□-Cox complex is strictly dependent on FixI (Fig. 2B).

To determine whether the presence of FixI is essential for the stability of the *aa_3_*_-_ and/or *cbb*□-Cox complex, we compared the proteomes of the WT and Δ*fixI* strains grown aerobically in PYE liquid medium. No significant difference in the levels of both *aa_3_* and *cbb*□-Cox subunits was observed between the two strains, indicating that the stability of *aa_3_* and *cbb*□-Coxs is not compromised by the absence of FixI (Fig. 2D).

The fact that *C. vibrioides* lacks a canonical CopA-type ATPase for Cu detoxification, but relies on FixI for *cbb*□-Cox function, led us to wonder whether this pattern was conserved across Alphaproteobacteria. Therefore, we analyzed 1,255 representative genomes with available Pfam annotations, focusing on the taxonomic distribution, domain architecture, and genomic context of Cu- transporting P-type ATPases, particularly their co-occurrence with *cbb*□-Cox gene clusters (Fig. 2E). Two distinct domain architectures were identified among these P-type ATPases: a 1-domain variant (PF00403–PF00122–PF00702) containing a single N-terminal heavy metal-binding domain (PF00403), and a 2-domain variant (PF00403–PF00403–PF00122–PF00702) featuring a duplication of this domain. The 1-domain variant frequently co-occurs with *cbb*□-Cox clusters in members of Caulobacterales, suggesting a functional association with FixI. Notably, Caulobacterales exhibit a marked reduction in the overall number of P-type ATPases compared to other Alphaproteobacterial orders, possibly reflecting adaptation to oligotrophic environments (Fig. 2E and Fig. S6).

In contrast, Rhodobacterales, Rhizobiales, and Sphingomonadales show greater diversity and higher copy numbers of both domain architectures, including the 2-domain variant, which may indicate functional diversification or redundancy. The consistent association of FixI with the 1-domain architecture supports its potential specialization for Cu delivery to the *cbb*□-Cox system, in contrast to CopA-like proteins that commonly contain duplicated metal-binding domains (Fig. 2E). Building on the genomic insights suggesting specialized Cu transport systems, we next examined the protein factors involved in Cu delivery essential for the full activity of *cbb*□- and *aa*□-Coxs.

### The full activity of *cbb***□**and *aa***□**-Cox enzymes requires a PCu_A_C Cu chaperone

Previous studies in *R. capsulatus* and *R. sphaeroides* have shown that the assembly of *cbb*□-Cox relies on the two periplasmic Cu chaperones PccA, a PCuAC-like protein, and SenC, a Sco-like protein (16, 19). One PCuAC homolog (hereafter referred to as PccA) and two Sco-like paralogs, Sco1 and Sco2, are encoded by the *C. vibrioides* genome. A multiple sequence alignment of PccA from *C. vibrioides* with the *R. capsulatus* and *R. sphaeroides* PccA reveals the presence of the canonical Cu-binding H(M)X□□MX□□HXM motif (Fig. S7A). Structural predictions using AlphaFold 3 indicate that PccA adopts a conserved cupredoxin-like fold, commonly observed in periplasmic Cu chaperones. This fold consists of two Met and two His residues forming a structural core with tetrahedral geometry, facilitating Cu¹□ binding between two antiparallel β-strands (Fig. S7B). Similarly, Sco1 and Sco2 harbor a conserved Cu-binding CXXXCP motif and a distant His residue commonly found in Sco-like proteins (28) (Fig. S8A). In addition, AlphaFold 3-based structural predictions of Sco1 and Sco2 highlight a thioredoxin-like fold found in other Sco homologs (28), comprising two parallel β-strands at the N-terminus, two antiparallel β-strands, and a unique α-helix in the C-terminal region (Fig. S8B). This *in silico* analysis suggests that PccA, Sco1, and Sco2 are potential Cu chaperones involved in Cu trafficking and delivery to Cox enzymes in *C. vibrioides*.

Interestingly, a Δ*pccA* mutant exhibited a NADI^slow^ phenotype, while the double Δ*sco1*Δ*sco2* mutant remained NADI^+^, indicating that PccA, but not Sco1 and Sco2, is required for *cbb*□-Cox and/or *aa*□- Cox activity (Fig. 3A). The NADI^slow^ phenotype of PccA is complemented by providing a plasmidic copy of *pccA*, indicating the absence of polar effects of *pccA* mutation (Fig. 3A). The triple Δ*pccA*Δ*sco1*Δ*sco2* mutant exhibited a similar phenotype to the Δ*pccA* mutant (Fig. 3A), reinforcing the idea that PccA is required for the activity of Cox in *C. vibrioides*. We hypothesized that the NADI^slow^ phenotype of the Δ*pccA* mutant results from a reduced activity of *cbb*□ and/or *aa*□-Cox. To test this hypothesis, two double Δ*pccA*Δ*aa*□ and Δ*pccA*Δ*cbb*□ mutants were generated. While the Δ*cbb*□ mutant exhibited a NADI^slow^ phenotype, the Δ*pccA*Δ*cbb*□ mutant showed no Cox activity (NADI□), suggesting that *pccA* deletion leads to a complete *aa*□-Cox inactivation. The Δ*pccA*Δ*aa*□ strain retained a NADI^slow^ phenotype, which contrasts with the NADI^+^ phenotype of the Δ*aa*□ mutant, implying that PccA is required for full *cbb*□-Cox activity on PYE solid medium.

**Figure 3.**
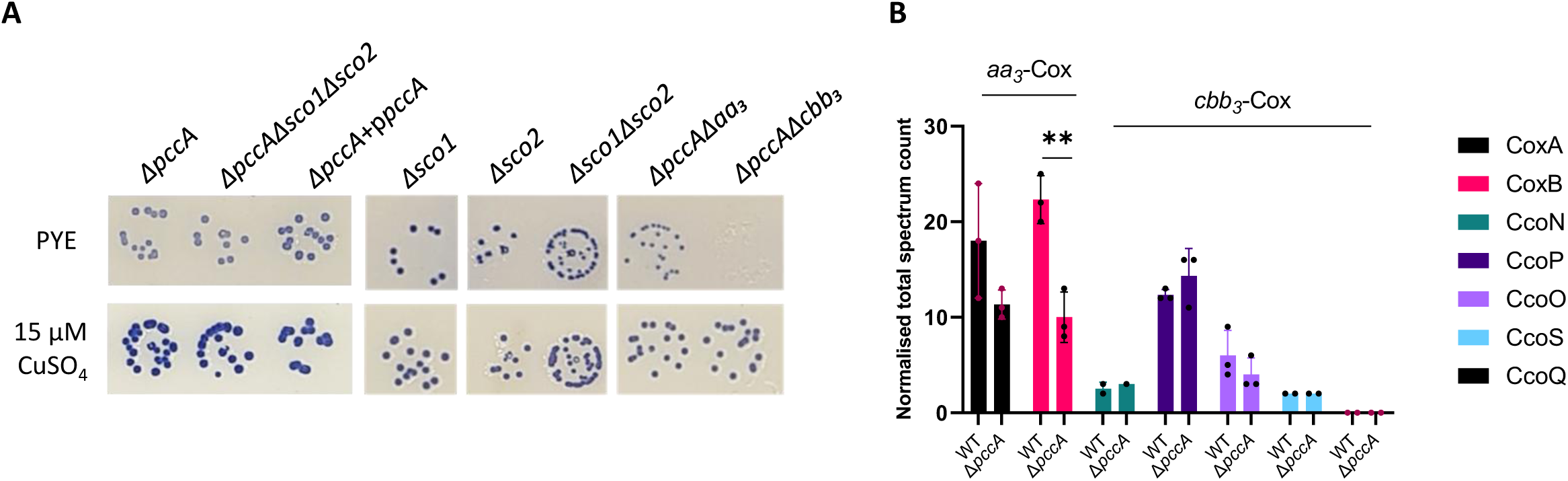
PccA is a Cu chaperone required for full *aa*□-Cox activity. **A.** Growth and NADI phenotypes of colonies of *C. vibrioides* WT and mutant strains. Cells were grown aerobically at 30°C on a PYE medium and PYE supplemented with 15 µM CuSO_4_, and the presence of Cox activity was visualized by NADI staining. Colonies that contain wild-type levels of Cox activity turn dark blue within a few seconds (NADI^+^), while those that have low or no Cox activity show lighter blue (NADI^slow^)or no blue staining (NADI^-^) upon longer exposure, respectively. Biological replicates = 3 (entire plate in Fig. S11B). **B.** Normalized total spectrum count of peptides of *aa*□-Cox subunits and *cbb*□-Cox subunits measured by LC-MS in both WT and Δ*pccA* strains grown in PYE medium. Individual values and means are represented. Biological replicates = 3. *P*-values were calculated using a t-test. (\**p* < 0.05, ** *p* < 0.01, *** *p* < 0.001 and \*\*\*\**p* < 0.0001) **(Table S4)**.

CuSO_4_ supplementation in the culture medium restored a WT Cox activity in the mutants lacking *pccA* (Δ*pccA*, Δ*pccA*Δ*sco1*Δ*sco2*, Δ*pccA*Δ*cbb*□ and Δ*pccA*Δ*aa*□), suggesting that Cu excess can compensate for the loss of PccA by ensuring Cu delivery to both *aa*□-Cox and *cbb*□-Cox (Fig. 3A). To assess the role of PccA in the stability of the *aa*□- and *cbb*□-Cox complexes, we performed a comparative LC-MS proteomic analysis of the WT and Δ*pccA* strains grown aerobically in a liquid PYE medium. The abundance of the *aa*□-Cox subunit CoxA was not significantly different between strains. However, the number of peptides of the *aa*□-Cox subunit CoxB was strongly reduced in the Δ*pccA* mutant, suggesting that PccA is necessary for *aa*□-Cox stability (Fig. 3B). On the other hand, the abundance of the *cbb*□-Cox subunits remained unchanged. None of the PccA, Sco1, and Sco2 chaperones appeared to play a role in Cu resistance in *C. vibrioides* (Additional information 2).

### A novel TonB-dependent receptor is specifically required for *aa***□**-Cox activity

Considering that FixI is dedicated to Cu delivery to *cbb*□-Cox only, we sought to find out how Cu was transported to *aa*□-Cox. Therefore, we performed a genetic screen on a mini-Tn5 transposon library realized in the Δ*fixI*Δ*cbb*□ genetic background, where *aa*□-Cox remains active (NADI^slow^), hypothesizing that transposon insertions in a gene essential for *aa*□-Cox activity (leading to a NADI□ phenotype) could reveal key players in this pathway. Several candidates were identified among the NADI^-^ mutants, with more than six mini-Tn5 insertions in the *CCNA_03615* gene encoding a TonB-dependent receptor (TBDR) referred to as TccA in this study (Table S1). To assess its potential role in *aa*□-Cox activity, we disrupted the *tccA* gene in the Δ*fixI*Δ*cbb*□ mutant, which exhibited minimal Cox activity (almost NADI□) (Fig. 4A), thereby validating the genetic screen. These data suggests that the TccA is required for the *aa*□-Cox activity. The loss of *aa_3_*-Cox activity was complemented by introducing an exogenous copy of *tccA* under the constitutive p*lac* promoter (Δ*fixI*Δ*cbb3,tccA::Tn5*+p*tccA*) using the pMR20 vector, ruling out any polar effect (Fig. 4B). *aa_3_*-Cox activity was fully restored in the triple Δ*fixI*Δ*cbb*□*,tccA::Tn5* mutant by supplementing with 15 µM CuSO_4_, but not with 25 µM FeSO_4_, suggesting that TccA is providing Cu ions to *aa*□-Cox (Fig. 4A). Consistent with this hypothesis, 3D structure prediction of TccA with AlphaFold 3 highlighted a potential Cu-binding site composed of the His116 and His321 residues (Fig. S10A).

**Figure 4.**
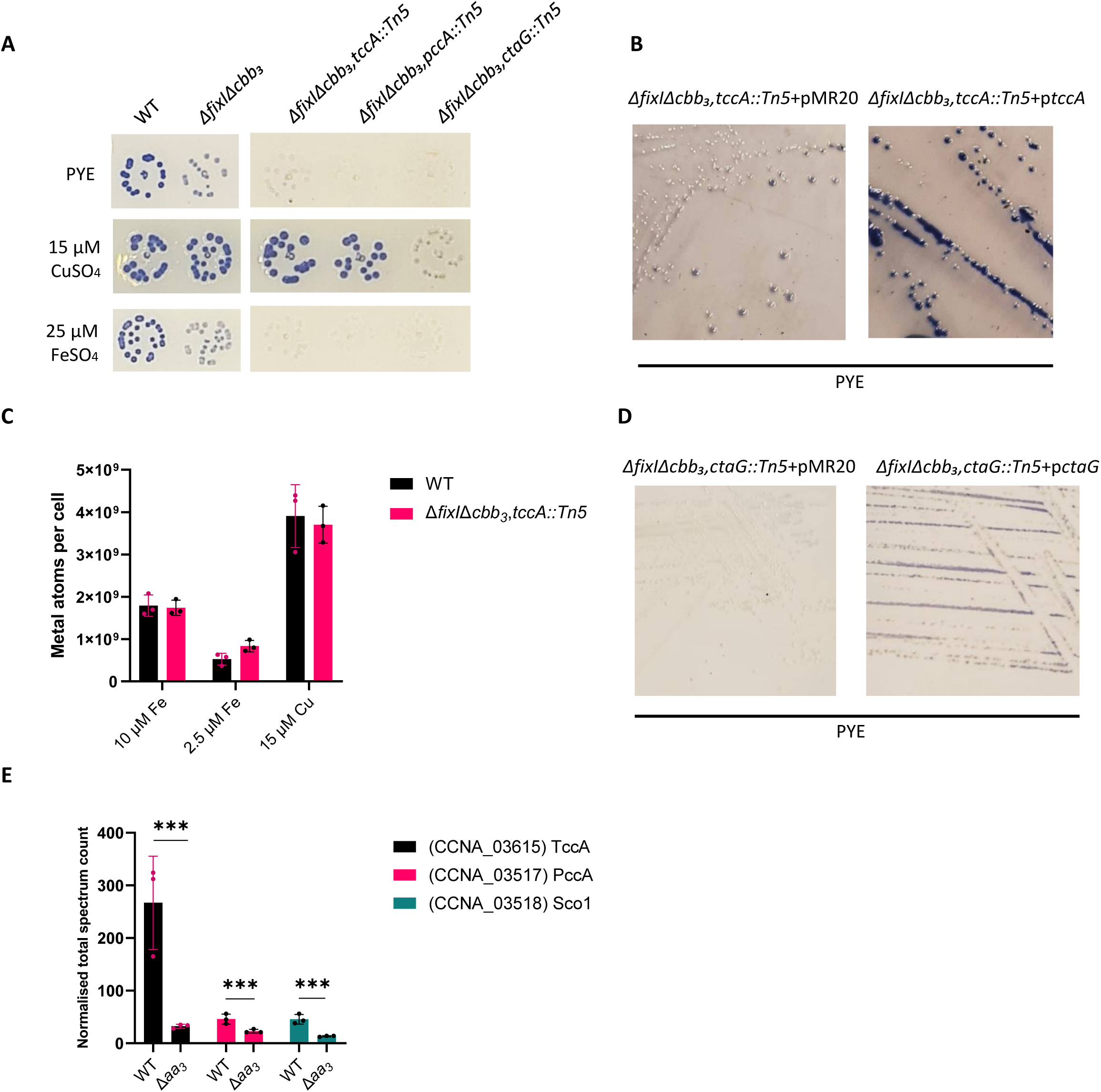
TccA is a TonB-dependent receptor required for the *aa*□-Cox activity. **A.** Growth and NADI phenotypes of colonies of *C. vibrioides* WT and mutant strains. Cells were grown aerobically at 30°C on a PYE medium, PYE supplemented with 15 µM CuSO_4_ and PYE supplemented with 25 µM FeSO_4_, and the presence of Cox activity was visualized by NADI staining. Colonies that contain WT levels of Cox activity turn dark blue within a few seconds (NADI^+^), while those that have low or no Cox activity show lighter blue (NADI^slow^) or no blue staining (NADI^-^) upon longer exposure, respectively. Biological replicates= 3. (entire plate in Fig. S12A) **B.** NADI phenotypes of colonies of *C. vibrioides* Δ*fixI*Δ*cbb*□*,tccA::Tn5* harboring empty pMR20 vector and Δ*fixI*Δ*cbb*□*,tccA::Tn5* strain carrying pMR20 vector with *tccA* gene. Cells were grown aerobically at 30°C on PYE medium supplemented with tetracycline to select the pMR20 vector. Biological replicates = 3. **C.** Fe and Cu atom counts per cell were measured in two independent experiments using cultures grown aerobically in minimal medium (M2G). In the first experiment, WT and Δ*fixI*Δ*cbb*□,*tccA::Tn5* strains were cultured overnight with 10 µM Fe and 2.5 µM Fe. In the second, WT and Δ*fixI*Δ*cbb*□*,tccA::Tn5* cultures were exposed for 30 min to 15 µM CuSO□. Data are shown as mean ± SD from at least three biological replicates. *P*-values were calculated using a t-test. (\**p* < 0.05, ** *p* < 0.01, *** *p* < 0.001 and \*\*\*\**p* < 0.0001) **(Table S4)**. **D.** NADI phenotypes of colonies of *C. vibrioides* Δ*fixI*Δ*cbb*□*,ctaG::Tn5* harboring empty pMR20 vector and Δ*fixI*Δ*cbb*□*,ctaG::Tn5* strain carrying pMR20 vector with *ctaG* gene. Cells were grown aerobically at 30°C on PYE medium supplemented with tetracycline to select the pMR20 vector. Biological replicates = 3. **E.** Normalized total spectrum count of peptides of *aa*□-Cox subunits and *cbb*□-Cox subunits in both WT and Δ*aa*□strain grown in PYE medium, measured by LC-MS. Individual values and means are represented. Biological replicates = 3. *P*-values were calculated using a t-test. (\**p* < 0.05, ** *p* < 0.01, *** *p* < 0.001 and \*\*\*\**p* < 0.0001) **(Table S4)**.

Given that TBDRs are typically involved in Fe transport, we measured intracellular Fe levels in the WT and Δ*fixI*Δ*cbb*□*,tbdr::Tn5* mutant using AAS. The cultures were aerobically grown in M2G minimal medium containing either 10 µM (optimal condition) and 2.5 µM (lower Fe condition) FeSO_4_.

No significant differences were observed in both conditions (Fig. 4B). Similarly, intracellular Cu levels remained unchanged between the WT strain and the Δ*fixI*Δ*cbb*□*tccA::Tn5* mutant when exposed for 30 min to moderate Cu stress (15 µM) in M2G (Fig. 4C). To determine whether TccA is exclusively dedicated to *aa*□-Cox, we constructed a double Δ*aa*□*,tccA::Tn5* mutant, where *cbb*□-Cox was still present. The mutant exhibited the same phenotype as the single Δ*aa*□ mutant, indicating that *cbb*□- Cox remains functional in the absence of TccA (Fig. S10B).

The genetic screen also identified the CtaG Cu chaperone, known to play a role in Cu delivery to the CuB center of *aa*□-Cox in *R. sphaeroides* (16). The Δ*fixI*Δ*cbb*□*,ctaG*::*Tn5* mutant exhibited a complete loss of *aa*□-Cox activity (NADI□) (Fig. 4A). Notably, Cu supplementation failed to rescue the phenotype, indicating that CtaG is essential for Cu delivery to *aa*□-Cox and, in turn, for its activity (Fig. 4A). The loss of *aa_3_*-Cox activity was partially complemented by introducing an exogenous copy of *ctaG* under the constitutive p*lac* promoter on the pMR20 vector (Δ*fixI*Δ*cbb3,tccA::Tn5*+p*ctaG*), confirming that the observed phenotype was not due to a polar effect (Fig. 4D). Interestingly, the abundance of TccA, PccA, and Sco1, which are encoded by neighboring genes, was reduced in the Δ*aa*□ mutant, suggesting a potential feedback loop between TccA, PccA, Sco1, and *aa*□-Cox. (Fig. 4E).

### *aa***□**-cox plays a key role under normoxic conditions

To determine whether the differential dominance of the *cbb*□- and *aa*□-Cox on solid media arises from O□ availability, and given that we cannot directly monitor enzyme levels on plates, we adapted the N,N,N′, N′-tetramethyl-p-phenylenediamine TMPD oxidase assay to *C. vibrioides* liquid cultures to functionally compare Cox activity under differing O□ conditions.

Interestingly, only the Δ*aa*□, Δ*pccA*, and Δ*pccA*Δ*sco1*Δ*sco2* mutants were impaired in Cox activity despite the presence of *cbb*□-Cox, confirming that PccA is essential for *aa*□-Cox activity in solid and liquid conditions (Fig. 5A and Fig. 5B). Besides, both Sco1 and Sco2 displayed no significant impact on the Cox activity (Fig. 5B), confirming the observed phenotype in solid medium.

**Figure 5.**
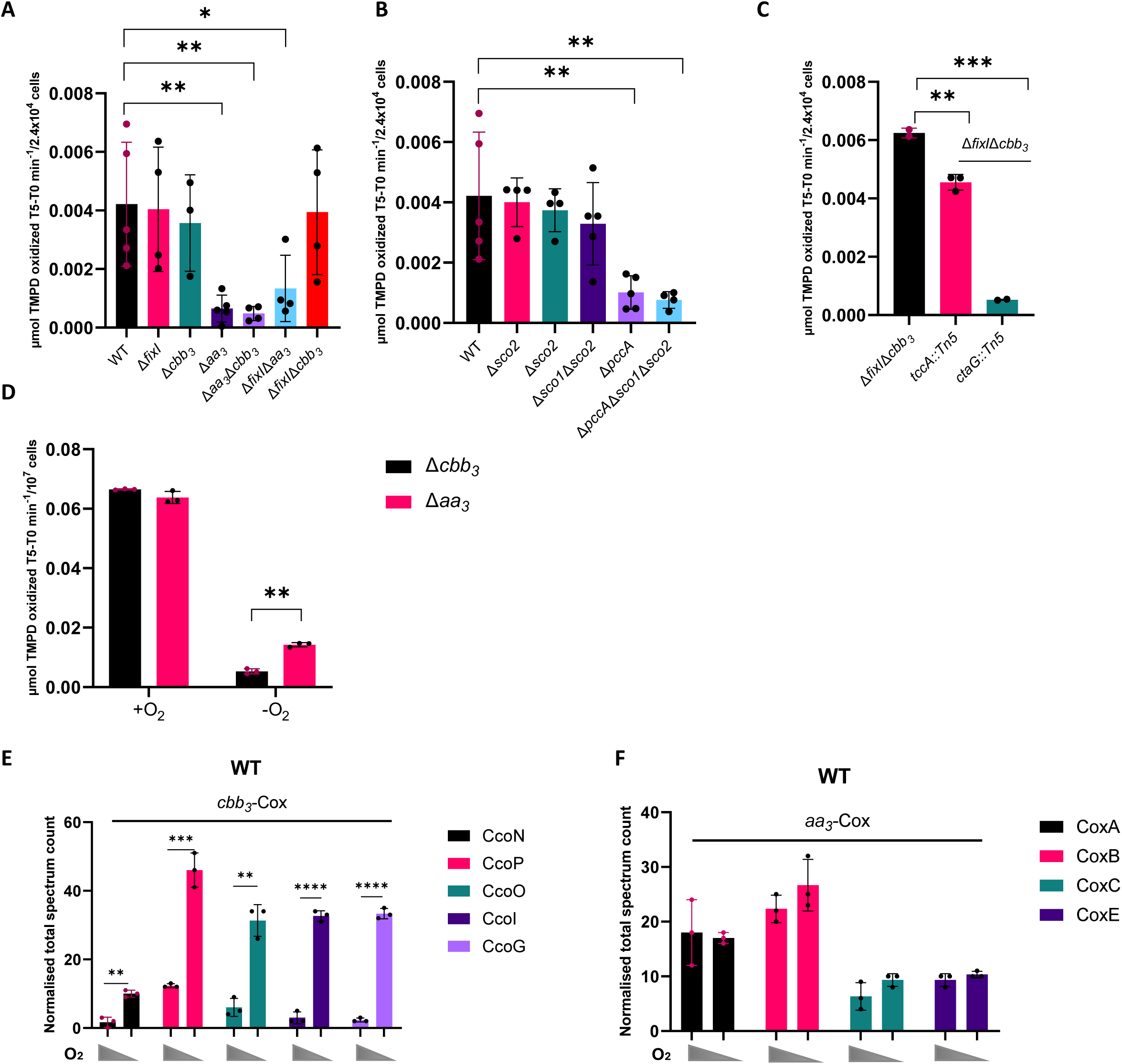
*cbb*□-Cox is more expressed and required under microaerobic conditions. **A, B, C.** TMPD oxidase activity in whole cells of *C. vibrioides* WT and mutant strains. Cells were cultured aerobically in a PYE-rich medium. Each value represents the mean of three independent cultures. Biological replicates = at least 2. *P*-values were calculated using a t-test. (\**p* < 0.05, ** *p* < 0.01, *** *p* < 0.001 and \*\*\*\**p* < 0.0001) **(Table S4)**. **D.** TMPD oxidase activity in whole cells of Δ*aa*□ and Δ*cbb*□ cultured aerobically and microaerobically in a PYE-rich medium. Biological replicates = 3. *P*-values were calculated using a t-test (\**p* < 0.05, ** *p* < 0.01, *** *p* < 0.001 and \*\*\*\**p* < 0.0001) **(Table S4)**. **E, F.** Normalized total spectrum count of peptides from *cbb*□-Cox and *aa*□-Cox subunits in the WT strain grown in PYE medium under aerobic and microaerobic conditions, as measured by LC-MS. Individual values and means are shown. Biological replicates = 3. *P*-values were calculated using a t-test (\**p* < 0.05, ** *p* < 0.01, *** *p* < 0.001 and \*\*\*\**p* < 0.0001) **(Table S4)**.

The Δ*fixI*Δ*cbb*□*,tccA::Tn5* mutant strain displayed an intermediate phenotype compared to the reference strain Δ*fixIcbb*□ (Fig. 5C). However, the Δ*fixI*Δ*cbb*□*,ctaG::Tn5* mutant strain showed no detectable Cox activity, indicating that CtaG is essential for *aa*□-Cox activity in liquid medium (Fig. 5C).

Subsequently, we generated microaerobic conditions in liquid medium and compared the Cox activity of the Δ*cbb*□ and Δ*aa*□ mutants using the TMPD oxidase assay. Notably, the Δ*aa*□ mutant, where *cbb*□-Cox remains intact, exhibited higher Cox activity under microaerobic conditions than under aerobic conditions, while no significant difference was observed in the Δ*cbb*□ mutant (Fig. 5D). This observation was reinforced by MS, showing an upregulation of *cbb*□-Cox subunits (CcoN, CcoO, CcoP) and CcoI/FixI and CcoG/FixG Cu reductase under microaerobic conditions (Fig. 5E). In contrast, the expression of the *aa*□-Cox subunits (CoxA, CoxB, CoxC, and CoxE) remained unchanged (Fig. 5F).

## Discussion

### C. vibrioides P1B1-type ATPase specifically delivers Cu to cbb□-type cytochrome c oxidase

As a strictly aerobic bacterium, *C. vibrioides* relies on Cu-dependent Cox enzymes for respiration (29), necessitating tight Cu homeostasis to avoid toxicity while ensuring proper enzyme metallation. Previous work from our lab revealed a bimodal Cu stress response: swarmer cells escape Cu-rich environments through chemotaxis, whereas stalked cells activate a Cu detoxification system (30). The outer membrane protein PcoB plays a key role by exporting Cu to prevent intracellular accumulation (31). However, the mechanism of Cu transport across the IM in *C. vibrioides* has remained unknown.

To address this question, we investigated how Cu is exported from the cytoplasm and identified a single P1B1-type ATPase, FixI, as a potential candidate. While P1B1-type ATPases are commonly associated with Cu detoxification in bacteria such as CopA from *E. coli* (3) requiring the cytoplasmic Cu-chaperone CopZ (5), our findings reveal that FixI in *C. vibrioides* is dispensable for Cu resistance (32). The absence of a CopZ homolog in *C. vibrioides* aligns with our findings. Besides FixI, *C. vibrioides* has a second P1B-type ATPase, ZctP, likely a P1B4-type ATPase involved in Zn homeostasis. Unlike *C. vibrioides*, some bacterial species such as *R. capsulatus* and *P. aeruginosa* possess multiple P1B1-type ATPases with distinct functions. CopA-like ATPases exhibit low Cu affinity but high efflux rates, supporting their role in Cu detoxification, whereas CcoI-like ATPases have high Cu affinity, consistent with their role in delivering Cu to *cbb*□-Cox, as shown in *P. aeruginosa* (9). *C. vibrioides* FixI appears to function as a CcoI-like P-type ATPase. Notably, under Cu excess, *cbb*□-Cox activity still relies on FixI, indicating that its function is not bypassed when Cu is more abundant and that the proper insertion of Cu in the CuB of *cbb3* relies on FixI. *C. vibrioides* lacks the CcoA Cu importer described in *R. capsulatus* (18), raising the question about how Cu is imported into the cytoplasm. The cytoplasmic Cu chaperone CopZ, essential for *cbb*□-Cox activity in *R. capsulatus* (33) is also absent in *C. vibrioides*. Despite the absence of CcoA and CopZ, FixI in *C. vibrioides* is still able to facilitate Cu delivery to the *cbb*□-Cox, suggesting that it may receive Cu either directly from the cytoplasmic pool or via an alternative chaperone or pathway that remains to be characterized (Fig. 6). As in *R. sphaeroides*, we found that in *C. vibrioides*, the absence of the *cbb*□- type Cox leads to a reduction in overall Cox activity. This remaining activity is attributed to the *aa*□- Cox, suggesting that *aa*□-Cox is functionally active but less efficient than *cbb*□-Cox on plates (34). In *C. vibrioides* mutants lacking *fixI* or *cbb*□, Cu supplementation restores Cox activity, suggesting Cu enhances *aa*□-Cox function by improving maturation or efficiency. This Cu-dependent increase differs from *R. sphaeroides*, where Cu addition does not boost *aa*□-Cox activity in the absence of *cbb*□-Cox (34). These differences suggest species-specific adaptations in Cu homeostasis and terminal Cox regulation, reflecting distinct ecological niches or metabolic strategies. Although FixI is essential for *cbb*□-Cox activity, its absence does not destabilize individual complex subunits. However, it may impair proper folding or assembly of *cbb*□-Cox, compromising its function. This is supported by the undetectable *cbb*□-Cox activity in the Δ*fixIaa*□ strain, even under Cu excess.

**Figure 6.**
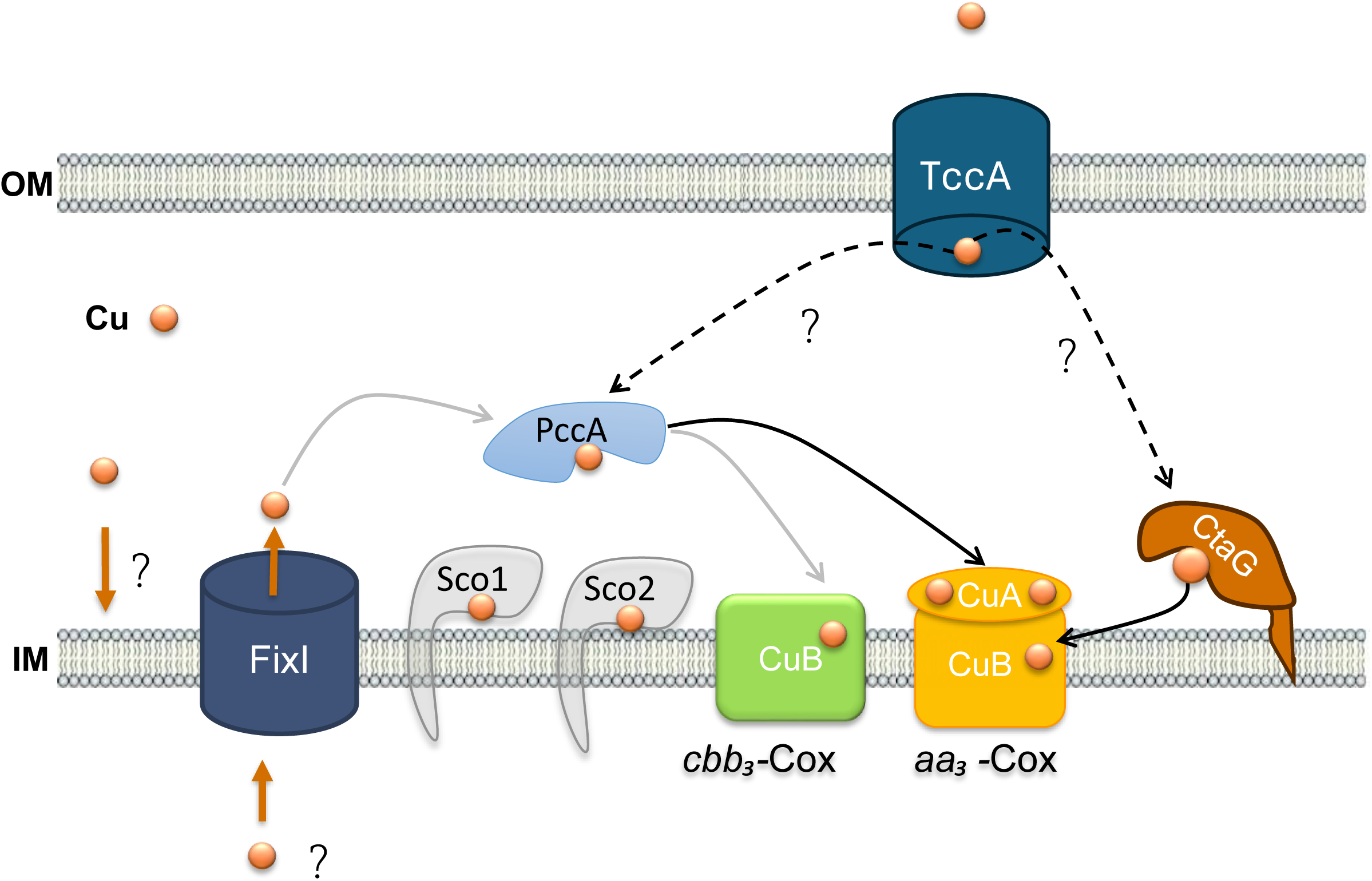
Putative Cu delivery pathway for *cbb*□ and *aa*□-Cox in *C. vibrioides*. ***aa*□-Cox route (in black):** Cu ions (orange spheres) are imported into the periplasm through a TonB-dependent receptor, TccA, in the outer membrane (OM). Once in the periplasm, PccA and CtaG bind Cu and deliver it to the CuA and CuB centers, respectively. A Cu exchange between PccA and CtaG is considered. ***cbb***□**-Cox route (in gray):** Cu is imported from the periplasm to the cytoplasm by a potential importer, FixI mediates the export, the exact transport mechanism, and the presence of a cytoplasmic Cu chaperone remain unknown. Once in the periplasm, Cu is delivered to the chaperone PccA, which probably interacts with Sco1 and ScoC2 to facilitate insertion of Cu into the CuB center of the catalytic CcoN subunit of *cbb*□-Cox. Together, these components ensure proper and safe incorporation of Cu into both CuA and CuB centers, which are essential for Cox function and respiratory activity in *C. vibrioides*. Solid arrows depict proven pathways or data, while dashed arrows represent speculative or unconfirmed interactions.

### PccA is solely essential for aa□-type Cox activity

Previous studies in *R. capsulatus* (19), *R.sphaeroides* (16), *B. japonicum* (20) and *Bacillus subtilis* (35) have demonstrated that PccA-like and/ or SenC/Sco-like proteins are essential for the Cu delivery pathway, facilitating correct Cu insertion into *aa*□ and *cbb_3_*-Cox (16, 36). In *R. sphaeroides*, the Sco-like protein is essential for assembling both the CuA and CuB centers in the *aa*□-type Cox, while the PccA-like protein assists in CuA center formation but has a secondary role (16). It likely functions upstream of the Sco-like protein in the Cu delivery pathway rather than acting as an independent chaperone (16). In contrast, *C. vibrioides* seems to rely solely on PccA, but not on Sco1 nor Sco2, to sustain *aa*□-Cox enzyme activity. Structural or mechanistic differences in the Cox complex of *C. vibrioides* may explain the divergence from typical Cu delivery pathways seen in other bacterial species. Previous studies have shown that PccA homologs from *Deinococcus radiodurans* and *C. vibrioides* bind Cu^1+^ through the metal-binding motif H(M)X□□MX□□HXM within a cupredoxin-like fold, suggesting a possible role in Cu delivery to the CuA center in Cox (36). Our results show a decreased abundance of the CoxB subunit, where CuA is located, in the absence of PccA, likely correlating with the lack of Cu bound to CuA. In Δ*pccAcbb*□ and Δ*pccAsco1sco2* strains, elevated Cu restores *aa*□-Cox activity, suggesting that under high Cu conditions, CuA can acquire Cu directly from the periplasm or indirectly via an unidentified Cu-binding chaperone.

### PccA is partially required for full cbb□-type Cox activity

The roles of PccA-like and Sco-like proteins in assembling the CuB center of the *cbb*□-Cox are closely related to their functions in CuA center assembly in the *aa*□-Cox in *R. sphaeroides*. Deletion of either leads to reduced CuB center assembly in the *cbb*□*-*type Cox, with Sco-like protein loss having a more substantial impact than that of the PccA-like protein (16). In contrast, in *B. japonicum*, only the PccA-like protein is essential for *cbb*□*-*Cox biogenesis, while the Sco-like protein does not appear to be required for CuB center assembly (20). In this study, Sco1 and Sco2 do not appear to be required for *cbb*□-type Cox activity, whereas PccA is essential for full function (Fig.6). This suggests that PccA delivers Cu to the CuB center of *cbb*□-type Cox. Supporting this, increased exogenous Cu compensates for the absence of PccA in the Δ*pccAaa*□ mutant and restores full *cbb*□-Cox activity. Although PccA is required for full *cbb*□-Cox activity, its absence does not affect the stability or accumulation of the complex’s subunits. This suggests that PccA operates at a later stage of assembly, likely mediating Cu insertion into the CuB catalytic center. In its absence, the complex may assemble but remains inactive.

### The activity of aa□-type Cox requires the presence of a specific TBDR

Several studies have investigated *cbb*□-specific Cu transporters, particularly in *R. capsulatus*, *R. sphaeroides*, and *P. aeruginosa*. No Cu transporter specific to *aa*□-type Cox was previously known. Our screen in *C. vibrioides* identified the outer membrane protein TccA as a likely missing component required for its maturation. In *B. japonicum*, Cu starvation induces a highly expressed operon encoding several periplasmic proteins, a TonB-dependent receptor (PcuB), and a cytoplasmic membrane protein (21). Among these, the PccA-like protein PcuC was required for CuA center formation in both aa□- and cbb□-Cox. In contrast, deletion of PcuB had no detectable phenotype, despite its predicted role as a Cu uptake receptor (21). Another key component identified in the genetic screen was the well-characterized Cox11-like protein, known as *ctaG*. Disruption of the *ctaG* gene completely abolished *aa*□-type Cox activity. Notably, external Cu supplementation could not bypass this defect, highlighting the essential role of CtaG in the maturation of the CuB site of *aa*□*-*Cox. This function has been previously demonstrated in *Saccharomyces cerevisiae* (37), *Paracoccus denitrificans*, and *R. sphaeroides* (11, 38, 39).

In *C. vibrioides*, we propose that Cu delivery to the *aa*□-type Cox involves a dedicated pathway composed of TccA, PccA, and CtaG. Cu is likely imported into the periplasm via TccA, which has a predicted His-rich Cu-binding site. In the periplasm, Cu is transiently bound by PccA for the insertion into the CuA center, while CtaG binds Cu for insertion into the CuB center of *aa*□-Cox. Additionally, Cu exchange may occur between PccA, CtaG, and possibly other chaperones, such as Sco1 and Sco2, forming a flexible and coordinated network. Disruption of any of the aforementioned key components abolishes *aa*□-Cox activity except Sco1 and Sco2 chaperones. However, since Cu supplementation restores activity in the absence of both PccA and TccA but not in the absence of CtaG, this suggests that Cu insertion into the CuA center can occur independently of PccA and TccA, while CuB site assembly remains strictly dependent on CtaG (Fig. 6). The reduced abundance of TccA and PccA in the absence of *aa*□-Cox suggests a feedback mechanism linking oxidase presence or activity to the expression or stability of its Cu-delivery components, preventing unnecessary production when the enzyme is absent.

### The expression and functional significance of both cbb□- and aa□-type Cox are O_2_-dependent

In this study, *aa*□*-Cox* was the primary oxidase under aerobic conditions and required PccA, CtaG, and TccA, whereas *cbb*□*-Cox* was upregulated under microaerobic conditions with increased subunit levels. Microarray data revealed that FixLJ, an O□-sensing two-component system, strongly activates *cbb*□*-Cox* transcription under low O□, indicating its primary role during O□ scarcity (29). For *aa*□*- Cox*, only minor expression changes occur under these conditions. The differences in *aa*□ and *cbb*□*- Cox* activities between solid and liquid media likely reflect O□ availability: in solid agar, slow O□ diffusion creates microaerobic or anaerobic zones deeper in colonies, while agitation in liquid cultures ensures better O□ distribution, favoring aerobic respiration and O□-dependent enzymes. Thus, *cbb*□*- Cox* predominates under microaerobic conditions, and *aa*□*-Cox* is essential in aerobic environments. Together, these findings reveal distinct Cu trafficking pathways supporting maturation of two terminal oxidases in *C. vibrioides* and identify TccA as a novel component specifically required for *aa*□*-Cox* biogenesis. The differential reliance on *cbb*□ and *aa*□*-Cox* under varying O□ conditions highlights a dynamic respiratory strategy and suggests greater diversity in mechanisms of bacterial Cu utilization.

### Experimental procedures

#### Bacterial strains, plasmids, and growth conditions

*C. vibrioides* NA1000 was routinely grown at 30°C under moderate shaking in PYE rich medium (0.2% bacto peptone, 0.1% yeast extract, 1 mM MgSO_4_, 0.5 mM CaCl_2_) or mineral M2G medium (5.4 mM Na_2_PO_4_, 4 mM KH_2_PO_4_, 9.35 mM NH_4_Cl, 0.2 % w/v 278 glucose, 10 μM FeSO_4_-EDTA, 0.5 mM MgSO_4_, 0.5 mM CaCl_2_). Microaerobic conditions were created by filling the 50 ml Falcon tube with PYE-rich medium and placing it in the incubator at 30°C without shaking. When required, the media were supplemented with 5 μg/mL kanamycin, 2.5 μg/mL oxytetracycline, 15 μg/mL nalidixic acid, and/or CuSO_4_. Exponentially growing cultures were used for all experiments. Strains and plasmids used in this study are listed in Table S2. The primers are provided in Table S3.

### Growth curve measurements

Bacterial cultures in the exponential growth phase (OD_660_ = 0.4–0.6) were diluted in PYE medium to a final OD660 nm of 0.05 and inoculated in 96-well plates with appropriate concentrations of CuSO4, ZnSO_4_, CdSO_4_, NiSO_4_, or CoSO_4_ when required. OD_660_ was recorded every 10 min for 24 h at 30°C under continuous shaking in an Epoch 2 absorbance reader (Biotek Instruments, Inc.).

### Viability assay

Overnight cultures in the stationary phase were diluted in 1:10 serial dilutions up to 10^−8^. Drops of 5 μL of each dilution were spotted on PYE or M2G plates containing BCS, CuSO_4_, ZnSO_4_, CdSO_4_, NiSO_4_, or CoSO_4_ at appropriate concentrations when required. The plates were incubated for 48 h at 30°C.

### Determination of metal content

Exponentially grown *C. vibrioides cultures* of 30 mL were fixed for 20 min in 2% paraformaldehyde (PFA) at 4°C. The bacteria were centrifuged for 10 min at 6,000 x g at 4°C and washed three times in 10 mL of ice-cold wash buffer [10 mM Tris–HCl (pH 6.8) and 100 μM EDTA]. Cells were resuspended in 2 mL MilliQ water and lyzed under 2.4 kbar by using a cell disrupter (Cell Disruption System, One-shot Model, Constant). Cell debris was removed by centrifugation at 10,000 x g for 15 min, and cell lysates were diluted in 1% HNO3. Samples were finally analyzed using an atomic absorption spectrophotometer (AA-7800) from Shimadzu. The number of bacteria for each sample was calculated based on the OD_660_ measurement taken prior to lysis. The ratio between the OD_660_ and the number of bacteria was determined in (40) Cellular metal concentrations were calculated using the following formula:

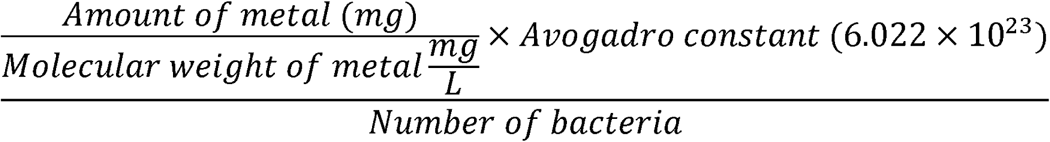

### Cytochrome *c* Oxidase Activity Assay

#### 1. Cox Activity on Solid Media

The activity of *cbb*□-type and *aa_3_*-type cytochrome *c* oxidase (Cox) was qualitatively assessed using the NADI staining reaction, which detects Cox activity based on the formation of indophenol blue. Plates containing colonies were flooded with a 1:1 (vol/vol) mixture of 35 mM α-naphthol and 30 mM *N*, *N*-dimethyl-*p*-phenylenediamine. Active colonies developed a dark blue coloration (NADI□) within 30 sec to 1 min. Colonies with reduced activity exhibited light blue staining (NADI^slow^) within 15 min, whereas colonies lacking both *aa_3_*and *cbb*□-Cox activity showed no color change (NADI□).

#### 2. Cox Activity in Liquid Media

Respiratory *c*-type cytochromes catalyze the oxidation of the artificial electron donor *N*, *N*, *N*′, *N*′- tetramethyl-p-phenylenediamine (TMPD), forming a blue-colored indophenol compound with a maximum absorbance at 520 nm (λ_max = 520 nm). Cox activity in whole cells was quantified spectrophotometrically by measuring the increase in absorbance at 520 nm at room temperature. At the late exponential phase, cultures grown in PYE were harvested, washed twice with 0.9% (w/v) NaCl, and resuspended to an OD_660_ ≈ of 0.2 in 33 mM potassium phosphate buffer (pH 7.0). A 100 µL aliquot (approximately 2.4x10^4^ cells) was mixed with 3.57 µL of 0.54 M TMPD in a 96-well plate, and a kinetic experiment was run for 10 min. Control samples contained TMPD. Cox activity was expressed as µmol TMPD oxidized min□¹ per 2.4x10^4^ cells, using an extinction coefficient (ε) of 6.1 mM□¹ cm□¹ for TMPD, the values at timepoint 5 min were subtracted from those at timepoint 0 min. Data represent the mean ± standard deviation of at least three biological replicates.

### Genetic screen using mini-Tn5

To identify genes that affect *aa_3_*-Cox activity, a transposon mutagenesis screen was performed in the Δ*fixI*Δ*cbb_3_* double knockout strain of *C. vibrioides*. Conjugation was carried out using *E. coli* donor cells harboring a plasmid-encoded mini-Tn5 transposon. Cells were spotted onto non-selective PYE plates and incubated for 6–8 h at 30□°C to allow for conjugation.

Following mating, colonies were selected and plated on PYE plates containing Kanamycin for transposon selection and counterselection against *E. coli* (Nalidixic acid). Colonies were then screened for Cox activity using NADI staining, which detects oxidase activity through the formation of a blue indophenol dye. While the Δ*fixI*Δ*cbb_3_* parental strain shows a weak light-blue staining due to residual *aa_3_*-Cox oxidase activity, colonies with a complete loss of staining (NADI^-^ or white phenotype) were selected for further analysis.

### LC–MS

Aerobically and microaerobically grown Cells from 15 mL cultures of *C. vibrioides* were isolated by centrifugation for 10 min at 8,000 rpm at 4°C and washed three times with ice-cold wash buffer (10 mM Tris–HCl pH 6.8, 100 μM EDTA). Normalized pellets were resuspended in 2 mL of complete EDTA-free protease inhibitor cocktail (Roche, Mannheim, Germany). Cells were lysed under 2.4 kbar by using a cell disrupter (Cell Disruption System, One-shot model, Constant). For proper dissolution of membrane proteins, lysates are incubated for 30 min with 0.01% SDS at room temperature. Cell debris was removed by centrifugation for 15 min at 19,000 x g. The samples were treated using the optimized filter-aided sample preparation protocol. Briefly, the samples were loaded onto Millipore Microcon 30 MRCFOR030 Ultracel PL-30 filtersfilters that have been rinsed and washed beforehand with 1% formic acid (FA) and 8 M urea buffer (8 M urea in 0.1 M Tris buffer at pH 8.5), respectively. The proteins on the filter were then exposed to a reducing agent (dithiothreitol) and alkylated with iodoacetamide. The proteins were then finally digested overnight with trypsin. The final step of digestion is to transfer proteins in 20 μL of 2% acetonitrile (ACN) and 0.1% FA in an injection vial for inverted phase chromatography. The digest was analyzed using nano-LC–ESI–MS/MS tims TOF Pro (Bruker, Billerica, MA, USA) coupled with UHPLC nanoElute (Bruker).

Peptides were separated by nanoUHPLC (nanoElute, Bruker) on a 75 μm ID, 25 cm C18 column with integrated CaptiveSpray insert (Aurora, IonOpticks, Melbourne) at a flowflow rate of 400 nL/min at 50°C. LC mobile phase A was water with 0.1% formic acid (v/v), and B was ACN with formic acid 0.1% (v/v). Samples were loaded directly on the analytical column at a constant pressure of 800 bars. The digest (1 μL) was injected, and the organic content of the mobile phase was increased linearly from 2% B to 15% in 22 min, 15% B to 35% in 38 min, and 35% B to 85% in 3 min. Data acquisition on the tims TOF Pro was performed using Hystar 5.1 and timsControl 2.0. tims TOF Pro data were acquired using 100 ms TIMS accumulation time and mobility (1/K0) range from 0.6 to 1.6 Vs/cm². Mass-spectrometric analysis was carried out using the parallel accumulation serial fragmentation (PASEF) acquisition method (31). One MS spectrum was followed by 10 PASEF MSMS spectra per total cycle of 1.1 s.

All MS/MS samples were analyzed using Mascot (Matrix Science, London, UK; version 2.8.1). Mascot was set up to search the *C. vibrioides* NA1000_190306 database from UniRef 100 and Contaminants_20190304 database, assuming the digestion enzyme trypsin. Mascot was searched with a fragment ion mass tolerance of 0.050 Da and a parent ion tolerance of 15 PPM. Carbamidomethyl of cysteine was specified in Mascot as a fixed modification. Oxidation of methionine and acetyl of the n-terminus was specified in Mascot as variable modifications.

Scaffold (version Scaffold_5.1.1,Scaffold_5.1.1, Proteome Software, Inc., Portland, OR) was used to validate MS/MS-based peptide and protein identifications. Peptide identifications were accepted if they could be established at a probability greater than 97.0% to achieve an FDR of less than 1.0% by the Percolator posterior error probability calculation (32). Protein identifications were accepted if they could be established at a probability greater than 50.0% to achieve an FDR of less than 1.0% and contained at least two identified peptides. Protein probabilities were assigned by the Protein Prophet algorithm (33). Proteins that contained similar peptides and could not be differentiated based on MS/MS analysis alone were grouped to satisfy the principles of parsimony. Proteins sharing significant peptide evidence were grouped into clusters.

### Statistical analysis

Statistical analyses were performed as required. All comparisons were analyzed using an unpaired t-test. A p-value below 0.05, 0.01, 0.001, and 0.0001 is indicated by *, **, ***, and ****, respectively (Table S4).

### Genome dataset and domain annotation

A set of bacterial proteomes from the UniProt Reference Proteomes dataset (UniProt Consortium, 2023) was annotated with Pfam domains using pfam_scan.pl and Pfam-A HMM profiles (Mistry et al., 2021). These annotated proteomes were then cross-referenced with assemblies from the Genome Taxonomy Database (GTDB, release R214; Parks et al., 2022), retaining only those genomes for which a matching GTDB assembly was available. This intersection resulted in a curated dataset of 1,255 representative Alphaproteobacterial genomes with both phylogenetic placement and domain annotations.

Identification of P-type ATPases with distinct domain architectures Proteins annotated as P-type ATPases were extracted based on the presence of specific Pfam domain architectures:

• **1-domain variant**: PF00403–PF00122–PF00702

• **2-domain variant**: PF00403–PF00403–PF00122–PF00702

These configurations differ by a duplication of the N-terminal heavy metal-binding domain (PF00403). Candidate proteins matching either variant were retrieved across the Alphaproteobacterial dataset and linked to their respective genomes.

### Extraction and analysis of genomic context

For each ATPase of interest, its genomic neighborhood was extracted by identifying all adjacent genes located on the same strand (i.e., transcribed in the same direction), both upstream and downstream of the ATPase gene. This strand-based criterion enabled the retrieval of full co-directional gene clusters, which were then annotated to identify potential functional elements, including cbb□-type cytochrome c oxidase subunits (e.g., FixC, FixP, FixO), regulatory elements, and additional neighboring genes.

Genomic contexts containing multiple *cbb*□-type subunits in proximity to a 1-domain ATPase were annotated as FixI-like clusters. ATPases lacking such co-localization were left unclassified in functional terms, as they may represent CopA-like or other metal-transporting ATPases with distinct roles.

### Phylogenetic mapping and visualization

The GTDB bacterial phylogeny was pruned to retain only the selected Alphaproteobacterial genomes. Mapping of ATPase presence, domain architecture, copy number, and associated functional classification onto the tree was performed using iTOL v6 (Letunic and Bork, 2021). Additional data visualizations were generated using custom Python scripts. The final figure summarizes the phylogenetic distribution of ATPase types, color-coded by copy number and co-occurrence with *cbb*□-type oxidase clusters (Figure 2E).

## Supporting information

This article contains supporting information

## Acknowledgements

We acknowledge the group team and the URBM members for fruitful discussions. We thank Rob Van Houdt for the information regarding the genome analysis. We thank the thesis committee members, Xavier De Bolle and Jean-François Collet, for their insightful comments.

## Data Availability

All data are contained within the manuscript and Supporting Information section. The raw data can be shared upon request to

## References

1. Argüello, J. M., Raimunda, D., and Padilla-Benavides, T. (2013) Mechanisms of copper homeostasis in bacteria. Front Cell Infect Microbiol. 3, 73

2. Andrei, A., Öztürk, Y., Khalfaoui-Hassani, B., Rauch, J., Marckmann, D., Trasnea, P. I., Daldal, F., and Koch, H. G. (2020) Cu homeostasis in bacteria: The ins and outs. Membranes (Basel). 10, 1– 45

3. Rensing, C., Fan, B., Sharma, R., Mitra, B., Rosen, B. P., and Kaback, H. R. CopA: An Escherichia coli Cu(I)-translocating P-type ATPase. 97, 652–656

4. Fan, B., and Rosen, B. P. (2002) Biochemical characterization of CopA, the Escherichia coli Cu(I)- translocating P-type ATPase. Journal of Biological Chemistry. 277, 46987–46992

5. Drees, S. L., Klinkert, B., Helling, S., Beyer, D. F., Marcus, K., Narberhaus, F., and Lübben, M. (2017) One gene, two proteins: coordinated production of a copper chaperone by differential transcript formation and translational frameshifting in Escherichia coli. Mol Microbiol. 106, 635–645

6. Multhaup, G., Strausak, D., Bissig, K. D., and Solioz, M. (2001) Interaction of the CopZ copper chaperone with the CopA copper ATPase of Enterococcus hirae assessed by surface plasmon resonance. Biochem Biophys Res Commun. 288, 172–177

7. Wimmer, R., Herrmann, T., Solioz, M., and Wüthrich, K. (1999) NMR structure and metal interactions of the CopZ copper chaperone. Journal of Biological Chemistry. 274, 22597–22603

8. Andrei, A., Di Renzo, M. A., Öztürk, Y., Meisner, A., Daum, N., Frank, F., Rauch, J., Daldal, F., Andrade, S. L. A., and Koch, H. G. (2021) The CopA2-Type P1B-Type ATPase CcoI Serves as Central Hub for cbb3-Type Cytochrome Oxidase Biogenesis. Front Microbiol. 12, 712465

9. González-Guerrero, M., Raimunda, D., Cheng, X., and Argüello, J. M. (2010) Distinct functional roles of homologous Cu+ efflux ATPases in Pseudomonas aeruginosa. Mol Microbiol. 78, 1246– 1258

10. Ekici, S., Pawlik, G., Lohmeyer, E., Koch, H. G., and Daldal, F. (2012) Biogenesis of cbb3-type cytochrome c oxidase in Rhodobacter capsulatus. Biochim Biophys Acta Bioenerg. 1817, 898– 910

11. Hederstedt, L. (2022) Diversity of Cytochrome c Oxidase Assembly Proteins in Bacteria. Microorganisms. 10, 926

12. Kawakami, T., Kuroki, M., Ishii, M., Igarashi, Y., and Arai, H. (2010) Differential expression of multiple terminal oxidases for aerobic respiration in Pseudomonas aeruginosa. Environ Microbiol. 12, 1399–1412

13. Kaila, V. R. I., and Wikström, M. (2021) Architecture of bacterial respiratory chains. Nat Rev Microbiol. 19, 319–330

14. Canonica, F., Hennecke, H., and Glockshuber, R. (2019) Biochemical pathway for the biosynthesis of the CuA center in bacterial cytochrome c oxidase. FEBS Lett. 593, 2977–2989

15. Khalfaoui-Hassani, B., Verissimo, A. F., Shroff, N. P., Ekici, S., Trasnea, P.-I., Utz, M., Koch, H.-G., and Daldal, F. (2016) Biogenesis of Cytochrome c Complexes: From Insertion of Redox Cofactors to Assembly of Different Subunits. 41, pp. 527–554

16. Thompson, A. K., Gray, J., Liu, A., and Hosler, J. P. (2012) The roles of Rhodobacter sphaeroides copper chaperones PCuAC and Sco (PrrC) in the assembly of the copper centers of the aa3-type and the cbb3-type cytochrome c oxidases. Biochim Biophys Acta Bioenerg. 1817, 955–964

17. Dash, B. P., Alles, M., Bundschuh, F. A., Richter, O. M. H., and Ludwig, B. (2015) Protein chaperones mediating copper insertion into the CuA site of the aa3-type cytochrome c oxidase of Paracoccus denitrificans. Biochim Biophys Acta Bioenerg. 1847, 202–211

18. Ekici, S., Yang, H., Koch, H. G., and Daldal, F. (2012) Novel transporter required for biogenesis of cbb3-type Cytochrome C oxidase in Rhodobacter capsulatus. mBio. 3, 1–11

19. Trasnea, P. I., Utz, M., Khalfaoui-Hassani, B., Lagies, S., Daldal, F., and Koch, H. G. (2016) Cooperation between two periplasmic copper chaperones is required for full activity of the cbb3-type cytochrome c oxidase and copper homeostasis in Rhodobacter capsulatus. Mol Microbiol. 100, 345–361

20. Bühler, D., Rossmann, R., Landolt, S., Balsiger, S., Fischer, H. M., and Hennecke, H. (2010) Disparate pathways for the biogenesis of cytochrome oxidases in Bradyrhizobium japonicum. Journal of Biological Chemistry. 285, 15704–15713

21. Serventi, F., Youard, Z. A., Murset, V., Huwiler, S., Bühler, D., Richter, M., Luchsinger, R., Fischer, H. M., Brogioli, R., Niederer, M., and Hennecke, H. (2012) Copper starvation-inducible protein for cytochrome oxidase biogenesis in Bradyrhizobium japonicum. Journal of Biological Chemistry. 287, 38812–38823

22. Maertens, L., Cherry, P., Tilquin, F., Van Houdt, R., and Matroule, J. Y. (2021) Environmental conditions modulate the transcriptomic response of both Caulobacter crescentus morphotypes to cu stress. Microorganisms. 9, 1116

23. Altschup, S. F., Gish, W., Miller, W., Myers, E. W., and Lipman, D. J. (1990) Basic Local Alignment Search Tool. J. Mol. Biol. 215, 403–410

24. Grønberg, C., Hu, Q., Mahato, D. R., Longhin, E., Salustros, N., Duelli, A., Lyu, P., Bågenholm, V., Eriksson, J., Rao, K. U., Henderson, D. I., Meloni, G., Andersson, M., Croll, T., Godaly, G., Wang, K., and Gourdon, P. (2021) Structure and ion-release mechanism of PIB-4-type ATPases. Elife. 11, e73124

25. Buchwalow, I., Boecker, W., and Tiemann, M. (2015) The contribution of Paul Ehrlich to histochemistry: a tribute on the occasion of the centenary of his death. Virchows Archiv. 466, 111–116

26. Jumper, J., Evans, R., Pritzel, A., Green, T., Figurnov, M., Ronneberger, O., Tunyasuvunakool, K., Bates, R., Žídek, A., Potapenko, A., Bridgland, A., Meyer, C., Kohl, S. A. A., Ballard, A. J., Cowie, A., Romera-Paredes, B., Nikolov, S., Jain, R., Adler, J., Back, T., Petersen, S., Reiman, D., Clancy, E., Zielinski, M., Steinegger, M., Pacholska, M., Berghammer, T., Bodenstein, S., Silver, D., Vinyals, O., Senior, A. W., Kavukcuoglu, K., Kohli, P., and Hassabis, D. (2021) Highly accurate protein structure prediction with AlphaFold. Nature. 596, 583–589

27. Gourdon, P., Liu, X. Y., Skjorringe, T., Morth, J. P., Møller, L. B., Pedersen, B. P., and Nissen, P. (2011) Crystal structure of a copper-transporting PIB-type ATPase. Nature. 475, 59–65

28. Balatri, E., Banci, L., Bertini, I., Cantini, F., and Ciofi-Baffoni, S. (2003) Solution structure of Sco1: A thioredoxin-like protein involved in cytochrome c oxidase assembly. Structure. 11, 1431–1443

29. Crosson, S., Mcgrath, P. T., Stephens, C., Mcadams, H. H., and Shapiro, L. (2005) Conserved modular design of an oxygen sensory signaling network with species-specific output. PNAS. 102, 8018–8023

30. Lawareé, E., Gillet, S., Louis, G., Tilquin, F., Le Blastier, S., Cambier, P., and Matroule, J. Y. (2016) Caulobacter crescentus intrinsic dimorphism provides a prompt bimodal response to copper stress. Nat Microbiol. 1, 16098

31. Hennaux, L., Kohchtali, A., Bâlon, H., Matroule, J. Y., Michaux, C., and Perpète, E. A. (2022) Refolding and biophysical characterization of the Caulobacter crescentus copper resistance protein, PcoB: An outer membrane protein containing an intrinsically disordered domain. Biochim Biophys Acta Biomembr. 1864, 184038

32. Manuel Gonzalez-Guerrero, and José M. Arguello (2008) Mechanism of Cu+-transporting ATPases: Soluble Cu+ chaperones directly transfer Cu+ to transmembrane transport sites. PNAS. 105, 5992–5997

33. Utz, M., Andrei, A., Milanov, M., Trasnea, P. I., Marckmann, D., Daldal, F., and Koch, H. G. (2019) The Cu chaperone CopZ is required for Cu homeostasis in Rhodobacter capsulatus and influences cytochrome cbb3 oxidase assembly. Mol Microbiol. 111, 764–783

34. Khalfaoui-Hassani, B., Wu, H., Blaby-Haas, C. E., Zhang, Y., Sandri, F., Verissimo, A. F., Koch, H. G., and Daldal, F. (2018) Widespread distribution and functional specificity of the copper importer CcoA: Distinct Cu uptake routes for bacterial cytochrome c oxidases. mBio. 9, e00065–18

35. Mattatall, N. R., Jazairi, J., and Hill, B. C. (2000) Characterization of ypmQ, an accessory protein required for the expression of cytochrome c oxidase in Bacillus subtilis. Journal of Biological Chemistry. 275, 28802–28809

36. Banci, L., Bertini, I., Ciofi-Baffoni, S., Katsari, E., Katsaros, N., Kubicek, K., and Mangani, S. (2005) A copper(I) protein possibly involved in the assembly of CuA center of bacterial cytochrome c oxidase. PNAS. 102, 3994–3999.

37. Carr, H. S., George, G. N., and Winge, D. R. (2002) Yeast Cox11, a protein essential for cytochrome c oxidase assembly, is a Cu(I)-binding protein. Journal of Biological Chemistry. 277, 31237–31242

38. Hiser, L., Di Valentin, M., Hamer, A. G., and Hosler, J. P. (2000) Cox11p is required for stable formation of the Cu(B) and magnesium centers of cytochrome c oxidase. Journal of Biological Chemistry. 275, 619–623

39. Hannappel, A., Bundschuh, F. A., Greiner, P., Alles, M., Werner, C., Richter, M., and Ludwig, B. (2011) Bacterial model systems for cytochrome c oxidase biogenesis. Indian J. Chem. 50A, 374– 382.

40. Louis, G., Cherry, P., Michaux, C., Rahuel-Clermont, S., Dieu, M., Tilquin, F., Maertens, L., Van Houdt, R., Renard, P., Perpete, E., and Matroule, J. Y. (2023) A cytoplasmic chemoreceptor and reactive oxygen species mediate bacterial chemotaxis to copper. Journal of Biological Chemistry. 299, 105207

41. Zielazinski, E. L., Cutsail, G. E., Hoffman, B. M., Stemmler, T. L., and Rosenzweig, A. C. (2012) Characterization of a cobalt-specific P1B-ATPase. Biochemistry. 51, 7891–7900

42. Raimunda, D., Long, J. E., Sassetti, C. M., and Argüello, J. M. (2012) Role in metal homeostasis of CtpD, a Co 2+ transporting P1B4-ATPase of Mycobacterium smegmatis. Mol Microbiol. 84, 1139–1149

43. Guan, G., Pinochet-Barros, A., Gaballa, A., Patel, S. J., Argüello, J. M., and Helmann, J. D. (2015) PfeT, a P1B4-type ATPase, effluxes ferrous iron and protects Bacillus subtilis against iron intoxication. Mol Microbiol. 98, 787–803

44. Tsirigos, K. D., Peters, C., Shu, N., Käll, L., and Elofsson, A. (2015) The TOPCONS web server for consensus prediction of membrane protein topology and signal peptides. Nucleic Acids Res. 43, W401–W407

45. Grønberg, C., Hu, Q., Mahato, D. R., Longhin, E., Salustros, N., Duelli, A., Lyu, P., Bågenholm, V., Eriksson, J., Rao, K. U., Henderson, D. I., Meloni, G., Andersson, M., Croll, T., Godaly, G., Wang, K., and Gourdon, P. (2021) Structure and ion-release mechanism of PIB-4-type ATPases. Elife. 11, e73124

46. Axelsen, K. B., and Palmgren, M. G. (1998) Evolution of substrate specificities in the P-type ATPase superfamily. J Mol Evol. 46, 84–101

47. Valencia, E. Y., Braz, V. S., Guzzo, C., and Marques, M. V (2013) Two RND proteins involved in heavy metal efflux in Caulobacter crescentus belong to separate clusters within proteobacteria. BMC Microbiol. 13, 79

