## Supporting tables for "Unraveling the pathway of Copper Delivery to Cytochrome *c* oxidases in the Free-Living Bacterium *Caulobacter vibrioides*"

**Table S1. Candidates from the genetic screen that were found to be required for *aa*$_3$-Cox activity in *C. vibrioides*.**

| Category | Candidates | Locus tag | Tn5 insertions |
|---|---|---|---|
| *aa3*-Cox | CoxA | CCNA_03518 | 7 |
| | CoxB | CCNA_03517 | 7 |
| | CoxC | CCNA_03513 | 1 |
| | hp | CCNA_03515 | 3 |
| *bc1* complex | ubiquinol-cytochrome *c* reductase iron-sulfur subunit | CCNA_00505 | 2 |
| | ubiquinol cytochrome *c* reductase *b* subunit | CCNA_00506 | 11 |
| | ubiquinol-cytochrome *c* reductase cytochrome *c1* subunit | CCNA_00507 | 4 |
| Cytochrome *c* maturation (Ccm) | Cytochrome *c*-type biogenesis protein (CcmH) | CCNA_02847 | 1 |
| | Cytochrome *c*-type biogenesis heme chaperone (CcmE) | CCNA_02849 | 2 |
| | Heme exporter protein C (CcmC) | CCNA_03786 | 1 |
| | Heme exporter protein A (CcmA) | CCNA_03782 | 1 |
| | disulfide interchange protein (CcmG) | CCNA_03788 | 2 |
| | Cytochrome *c* | CCNA_03030 | 2 |
| Heme A | Heme A synthase CtaA | CCNA_01438 | 1 |
| Thiol | disulfide interchange protein TlpA | CCNA_02293 | 2 |
| Chaperones | PccA | CCNA_03617 | 1 |
| | CtaG | CCNA_03514 | 2 |
| Transporter | TccA | CCNA_03615 | 7 |

**Table S2. Bacterial strains and plasmids used in this study**

| Strains | Genotype and/or phenotype | Reference or source |
|---|---|---|
| ***Caulobacter vibrioides*** | | |
| WT | Wild-type NA1000, synchronizable variant of CB15 | (37) |
| Δ*fixI* | Knock-out strain for *fixI* gene (CCNA_01473) | This study |
| Δ$cbb_3$ | Knock-out strain for the *ccoN*, *ccoO*, *ccoQ*, *ccoP* genes (CCNA_01467) (CCNA_01468) (CCNA_01469) (CCNA_01470) | This study |
| Δ$aa_3$ | Knock-out strain for the *coxB* and *coxA* genes (CCNA_03517), (CCNA_03518) | This study |
| Δ$aa_3$Δ$cbb_3$ | Double knock-out strain of both $aa_3$ and $cbb_3$ genes, as mentioned earlier | This study |
| Δ*fixI*Δ$cbb_3$ | Double knock-out strain of both *fixI* and $cbb_3$ genes | This study |
| Δ*fixI*Δ$aa_3$ | Double knock-out strain of both *fixI* and $aa_3$ genes | This study |
| Δ*sco1* | Knock-out strain for *sco1* gene (CCNA_03618) | This study |
| Δ*sco2* | Knock-out strain for *sco2* gene (CCNA_00249) | This study |
| Δ*sco1*Δ*sco2* | Double knock-out strain of both *sco1* and *sco2* genes | This study |
| Δ*pccA* | Knock-out strain for *pccA* gene (CCNA_03617) | This study |
| Δ*pccA*Δ*sco1*Δ*sco2* | Triple knock-out strain of the three genes *pccA*, *sco1* and *sco2* | This study |
| Δ*pccA*Δ$aa_3$ | Double knock-out strain of both *pccA* and $aa_3$ genes | This study |
| Δ*pccA*Δ$cbb_3$ | Double knock-out strain of both *pccA* and $cbb_3$ genes | This study |
| *fixI*: C398A-C400A | WT strain with the two-point C398A and C400A substitutions in the chromosomal *fixI* | This study |
| Δ*fixI*+p*fixI* | Knock-out strain for *fixI* gene carrying a copy of *fixI* on the pMR10 under the control of the plac promoter; KanR | This study |
| Δ*pccA*+p*pccA* | Knock-out strain for *pccA* gene carrying a copy of *pccA* on the pMR10 under the control of the lac promoter; KanR | This study |
| Δ*fixI*Δ$cbb_3$,*tccA::Tn5* | Double knock-out strain of both *fixI* and $cbb_3$ genes with a mini-Tn5 transposon inserted in the CCNA_03615 gene | This study |
| Δ*fixI*Δ$cbb_3$,*tccA::Tn5*+p*tccA* | Double knock-out strain of both *fixI* and $cbb_3$ genes with a mini-Tn5 transposon inserted in the CCNA_03615 gene carrying a copy of *tccA* on the pMR20 under the control of the plac promoter; TetR | This study |
| Δ*fixI*Δ$cbb_3$,*ctaG::Tn5* | Double knock-out strain of both *fixI* and $cbb_3$ genes with a mini-Tn5 transposon inserted in the CCNA_03514 gene | This study |
| Δ*fixI*Δ$cbb_3$,*ctaG::Tn5*+p*ctaG* | Double knock-out strain of both *fixI* and $cbb_3$ genes with a mini-Tn5 transposon inserted in the CCNA_03514 gene carrying a copy of *ctaG* on the pMR20 under the control of the plac promoter; TetR | This study |
| Δ*zctP* | Knock-out strain for *zctP* gene (CCNA_02811) | This study |
| Δ*zctP*Δ*fixI* | Double Knock-out strain for both *fixI* and *zctP* genes | This study |
| Δ*zctP*+p*zctP* | Knock-out strain for *zctP* gene carrying a copy of *zctP* on the pMR10 under the control of the lac promoter; KanR | This study |
| WT+ p*zctP* | Knock-out strain carrying a copy of *zctP* on the pMR10 under the control of the lac promoter; KanR | This study |
| **Plasmids** | | |
| pNTPS | mobRP4+ ori-R6K sacB; integrative vector in *C. vibrioides* for in-frame deletions; KanR | |
| pMR10 | Low copy and replicative vector in *C. vibrioides*; KanR | |
| pMR20 | Derivative of pMR10; TetR | |

**Table S3. Primers used in this study**

| Primer | Target | Sequence |
|---|---|---|
| Pr1 | Δ*fixI* (Upstream *fixI* FWD) | AATGACGATCGCCGCGCAAT |
| Pr2 | Δ*fixI* (Upstream *fixI* overlap region RWD) | CGAACATCGCCGCCgATCTTGCTGATGCAGCCGGCGC |
| Pr3 | Δ*fixI* (Downstream *fixI* overlap region FWD) | GGCTGCATCAGCAAGATcGGCGGCGATGTTCGGAC |
| Pr4 | Δ*fixI* (Downstream *fixI* RWD) | GGCTTGATGTCATTGCCGACA |
| Pr5 | Δ*cbb3* (Upstream *ccoN* FWD) | GCATCTGCGCTGAACCGCCT |
| Pr6 | Δ*cbb3* (Upstream *ccoN* overlap region RWD) | GCGTGGGCATGGGCCACGGACTCCCTGGGGCGAA |
| Pr7 | Δ*cbb3* (Downstream *ccoP* overlap region FWD) | CCCCAGGGAGTCCGTGGCCCATGCCCACGCTCATC |
| Pr8 | Δ*cbb3* (Downstream *ccoP* FWD) | CAGGCGCATCCGGGCGTTA |
| Pr9 | Δ*aa3* (Upstream *coxB* FWD) | CATATCGATGTCCCTCGTGA |
| Pr10 | Δ*aa3* (Upstream *coxB* overlap region RWD) | GGACGACGGTTCACCATCCCTCACGG |
| Pr11 | Δ*aa3* (Downstream *coxA* overlap region FWD) | CGGGCCGTGAGGGATGGTGAACCGTC |
| Pr12 | Δ*aa3* (Downstream *coxA* overlap region RWD) | GGCCAGCCAATTGACCGCAA |
| Pr13 | Δ*sco1* (Upstream *sco1* FWD) | CCTGCGCCGGCTGATCACGC |
| Pr14 | Δ*sco1* (Upstream *sco1* overlap region RWD) | AAGGATTTGCCCGGGCAGTGATGCATCCCCGACAT |
| Pr15 | Δ*sco1* (Downstream *sco1* overlap region FWD) | GGGGATGCATCACTGCCCGGGCAAATCCTTTCCCG |
| Pr16 | Δ*sco1* (Downstream *sco1* RWD) | GGGATCTGCGTGCGCGCGT |
| Pr18 | Δ*sco2* (Upstream *sco2* FWD) | CAGGGTGGCCGCTGCGAGGG |
| Pr19 | Δ*sco2* (Upstream *sco2* overlap region RWD) | AATTTTTACGCCGGAGCGCGGTTCATCCTTTTACT |
| Pr20 | Δ*sco2* (Downstream *sco2* overlap region FWD) | AAGGATGAACCGCGCTCCGGCGTAAAAATTCGCGA |
| Pr21 | Δ*sco2* (Downstream *sco2* RWD) | GATGAAGACGGCGTGGTCGG |
| Pr22 | Δ*pccA* (Upstream *pccA* FWD) | TGACAGGCGGCTTCACCCTG |
| Pr23 | Δ*pccA* (Upstream *pccA* overlap region RWD) | GAGCGATGGCCCGCACTTCAGGCTCCGGTCTGTGC |
| Pr24 | Δ*pccA* (Downstream *pccA* overlap region FWD) | GACCGGAGCCTGAAGTGCGGGCCATCGCTCTGAC |
| Pr25 | Δ*pccA* (Downstream *pccA* RWD) | CCCCTTGGCGTCGAACAGAT |

| Primer | Target | Sequence |
|---|---|---|
| Pr26 | *fixI*: C398A-C400A (Upstream region FWD) | GGACAACGCCCTGATCACCG |
| Pr27 | *fixI*: C398A-C400A (Upstream overlap region RWD) | ACCGCCAGGCCCAGGGCTGCCGGTGCAGTCACGAT |
| Pr28 | *fixI*: C398A-C400A (Downstream overlap region FWD) | TGACTGCACCGGCAGCCCTGGGCCTGGCGGTGC |
| Pr29 | *fixI*: C398A-C400A (Downstream region RWD) | CAGCCGTCTCGACGCAGTCCT |
| Pr30 | Δ*zctP* (Upstream *zctP* FWD) | TGTTCGTCCTGCCGGCGAT |
| Pr31 | Δ*zctP* (Upstream *zctP* overlap region RWD) | GCGCGAGGTCGTTCGCGGCGCGTCTCCAGAGTTAG |
| Pr32 | Δ*zctP* (Downstream *zctP* overlap region FWD) | AACTCTGGAGACGCGCCGCGAACGACCTCGCGCCAA |
| Pr33 | Δ*zctP* (Downstream *zctP* RWD) | TCGTTCCTTCGTCACTTCGATG |
| Pr34 | Forward amplification of *zctP* (CCNA_02811) gene with restriction enzyme Xbal | tgctctagaATGGCCGCCGGCATAAAGCCC |
| P35 | Reverse amplification of *zctP* (CCNA_02811) gene with restriction enzyme KpnI | ggtaccTTCGCTCACGCCGTCCGGTC |
| Pr36 | Forward amplification of the *pccA* gene CCNA_03617 with restriction enzyme HindIII | CCCAAGCTTATGAAGACCCTGACCCTGCT |
| Pr37 | Reverse amplification of the *pccA* gene CCNA_03617 with restriction enzyme XbaI | TGCTCTAGATCAGTGATGCATCCCCGACA |
| Pr38 | Forward amplification of the *fixI* gene CCNA_01473 with restriction enzyme SacI | CCGAGCTCATGAGCCACAGCCTCGCC |
| Pr39 | Reverse amplification of the *fixI* gene CCNA_01473 with restriction enzyme XbaI | TGCTCTAGATCATCGGTTGAAACTCCG |
| Pr40 | Forward amplification of the *tccA* gene CCNA_03615 with restriction enzyme XbaI | TGCTCTAGAATGAAGTCTCTTCTGCTG |
| Pr41 | Reverse amplification of the *tccA* gene CCNA_03615 with restriction enzyme EcoRI | CCGGAATTCTTAGAAGCTGACGGTCA |
| Pr42 | Forward amplification of the *ctaG* gene CCNA_03513 with restriction enzyme HindIII | CCCAAGCTTATGTCGCAAAACCCCAC |
| Pr43 | Reverse amplification of the *ctaG* gene CCNA_03513 with restriction enzyme EcoRI | CCGGAATTCCTATAGACCTCTCGACGG |

**Table S4. p-values. A t-test was performed using GraphPad Prism.**

| Fig. 1D Unpaired t-test | Summary | Adjusted P Value |
|---|---|---|
| WT vs. Δ*fixI* | ns | 0.3452 |
| **Fig. 2D Unpaired t-test** | | |
| WT vs. Δ*fixI* (CoxA) | ns | 0.3503 |
| WT vs. Δ*fixI* (CoxB) | ns | 0.6520 |
| WT vs. Δ*fixI* (CoxC) | ns | 0.6240 |
| WT vs. Δ*fixI* (CcoN) | ns | 0.3118 |
| WT vs. Δ*fixI* (CcoP) | ns | 1 |
| WT vs. Δ*fixI* (CcoO) | ns | 0.3552 |
| WT vs. Δ*fixI* (CcoG) | ns | 0.08 |
| **Fig. 3B Unpaired t-test** | | |
| WT vs. Δ*pccA* (CoxA) | ns | 0.1356 |
| WT vs. Δ*pccA* (CoxB) | ** | 0.0043 |
| WT vs. Δ*pccA* (CcoN) | ns | 0.3118 |
| WT vs. Δ*pccA* (CcoP) | ns | 0.3046 |
| WT vs. Δ*pccA* (CcoO) | ns | 0.3349 |
| WT vs. Δ*pccA* (CcoS) | ns | 0.6779 |
| **Fig. 4C Unpaired t-test** | | |
| WT vs. Δ*fixIcbb3,tccA::Tn5* (10 mM Fe) | ns | 0.7889 |
| WT vs. Δ*fixIcbb3,tccA::Tn5* (2.5 mM Fe) | ns | 0.05125 |
| WT vs. Δ*fixIcbb3,tccA::Tn5* (15 µM Cu) | ns | 0.70344 |
| **Fig. 4E Unpaired t-test** | | |
| WT vs. Δ*aa3* (TccA CCNA_03615) | *** | 0.0101 |
| WT vs. Δ*aa3* (PccA CCNA_03617) | *** | 0.017 |
| WT vs. Δ*aa3* (Sco1 CCNA_03618) | *** | 0.0039 |

| **Fig. 5A Unpaired t-test** | **Summary** | **Adjusted P Value** |
|---|---|---|
| WT vs. Δ*fixI* | ns | 0.5283 |
| WT vs. Δ*cbb3* | ns | 0.1740 |
| WT vs. Δ*aa3* | ** | 0.0062 |
| WT vs. Δ*aa3*Δ*cbb3* | ** | 0.0039 |
| WT vs. Δ*fixI*Δ*aa3* | * | 0.019 |
| WT vs. Δ*fixI*Δ*cbb3* | ns | 0.4942 |
| **Fig. 5B Unpaired t-test** | | |
| WT vs. Δ*sco1* | ns | 0.409 |
| WT vs. Δ*sco2* | ns | 0.658 |
| WT vs. Δ*sco1*Δ*sco2* | ns | 0.5 |
| WT vs. Δ*pccA* | ** | 0.0023 |
| WT vs. Δ*pccA*Δ*sco1*Δ*sco2* | ** | 0.0052 |
| **Fig. 5B Unpaired t-test** | | |
| Δ*fixI*Δ*cbb3* vs. Δ*fixI*Δ*cbb3,tccA::Tn5* | ** | 0.0045 |
| Δ*fixI*Δ*cbb3* vs. Δ*fixI*Δ*cbb3,ctaG::Tn5* | *** | 0.0004 |
| **Fig. 5D Unpaired t-test** | | |
| Δ*cbb3* vs. Δ*aa3* (+O2) | ns | 0.1454 |
| Δ*cbb3* vs. Δ*aa3* (-O2) | ** | 0.0013 |
| **Fig. 5E Unpaired t-test** | | |
| CcoN (+O2 vs. -O2) | ** | 0.0014 |
| CcoP (+O2 vs. -O2) | *** | 0.0003 |
| CcoO (+O2 vs. -O2) | ** | 0.0012 |
| CcoI (+O2 vs. -O2) | **** | 0.0001 |
| CcoG (+O2 vs. -O2) | **** | 0.0001 |

| **Fig. 5F Unpaired t-test** | **Summary** | **Adjusted P Value** |
|---|---|---|
| CoxA (+O2 vs. -O2) | ns | 0.7900 |
| CoxB (+O2 vs. -O2) | ns | 0.2336 |
| CoxC (+O2 vs. -O2) | ns | 0.1338 |
| CoxE (+O2 vs. -O2) | ns | 0.2508 |
| **Fig. S4B Unpaired t-test** | | |
| WT vs. Δ*zctP* (PYE) | ns | 0.46 |
| WT vs. Δ*zctP* (Zn) | ns | 0.34 |
| **Fig. S5C Unpaired t-test** | | |
| WT vs. Δ*aa3*Δ*cbb3* (qoxA) | ns | 0.15 |
| WT vs. Δ*aa3*Δ*cbb3* (qoxB) | ns | 0.47 |
| WT vs. Δ*aa3*Δ*cbb3* (Surf1) | ns | 0.37 |
| WT vs. Δ*aa3*Δ*cbb3* (CydA) | ns | 0.41 |
| WT vs. Δ*aa3*Δ*cbb3* (CydB) | ns | 0.07 |
| WT vs. Δ*aa3*Δ*cbb3* (CydC) | ns | 0.44 |
| WT vs. Δ*aa3*Δ*cbb3* (CydD) | * | 0.04 |
