## Supporting figures for "Unraveling the pathway of Copper Delivery to Cytochrome *c* oxidases in the Free-Living Bacterium *Caulobacter vibrioides*"

### Figure S1

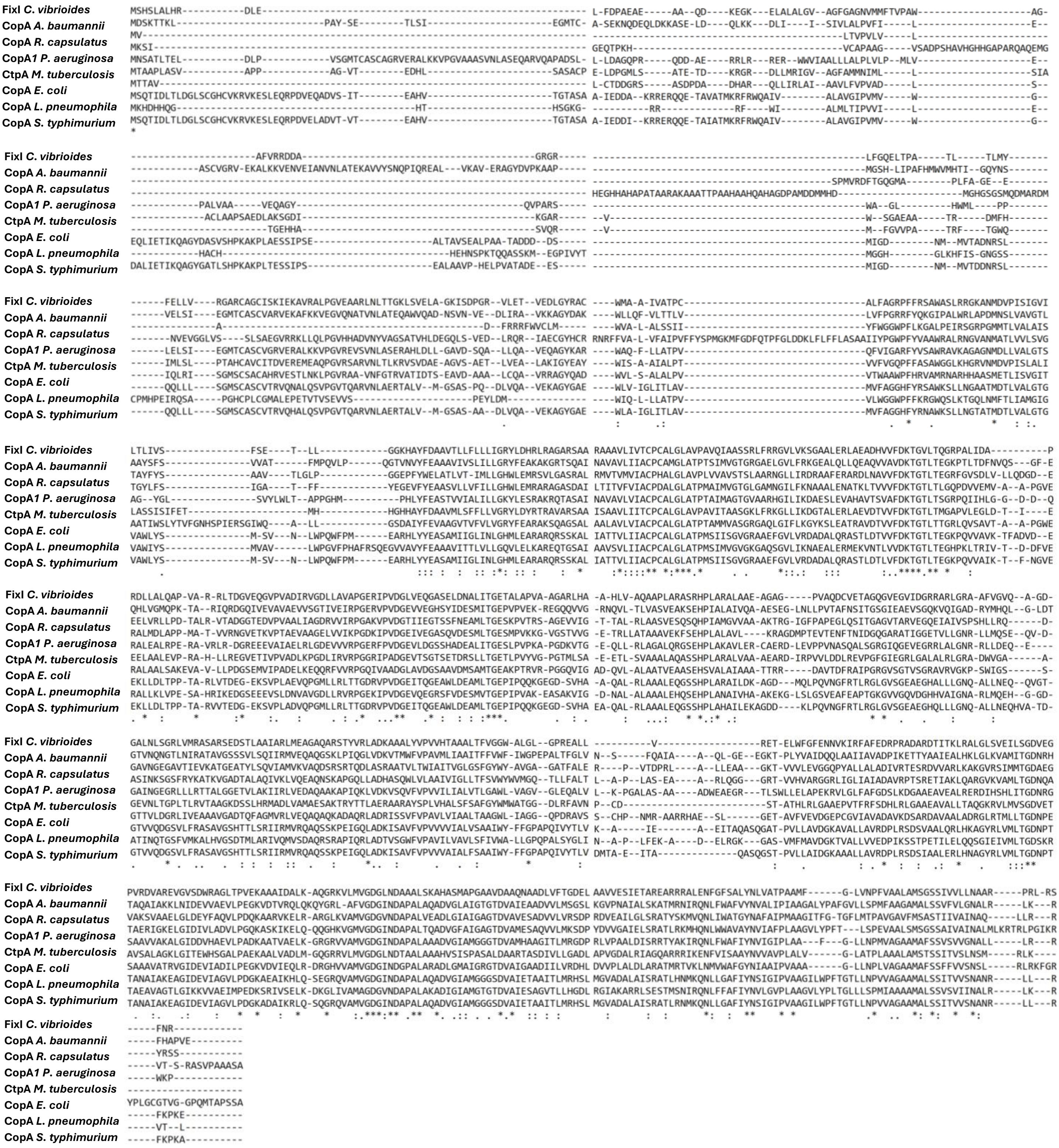


**Figure S1. Multiple sequence alignment of FixI from *C. vibrioides* and homologous P1B-type ATPases from diverse bacterial Species.** Whole amino acid sequence alignment of FixI from *C. vibrioides* with its homologs CopA from *Acinetobacter baumannii*, CopA from *Rhodobacter capsulatus*, CopA *Pseudomonas aeruginosa*, CtpA *Mycobacterium tuberculosis,* CopA *Escherichia coli*, CopA *Legionella pneumophila,* and CopA *Salmonella typhimurium*.

### Figure S2

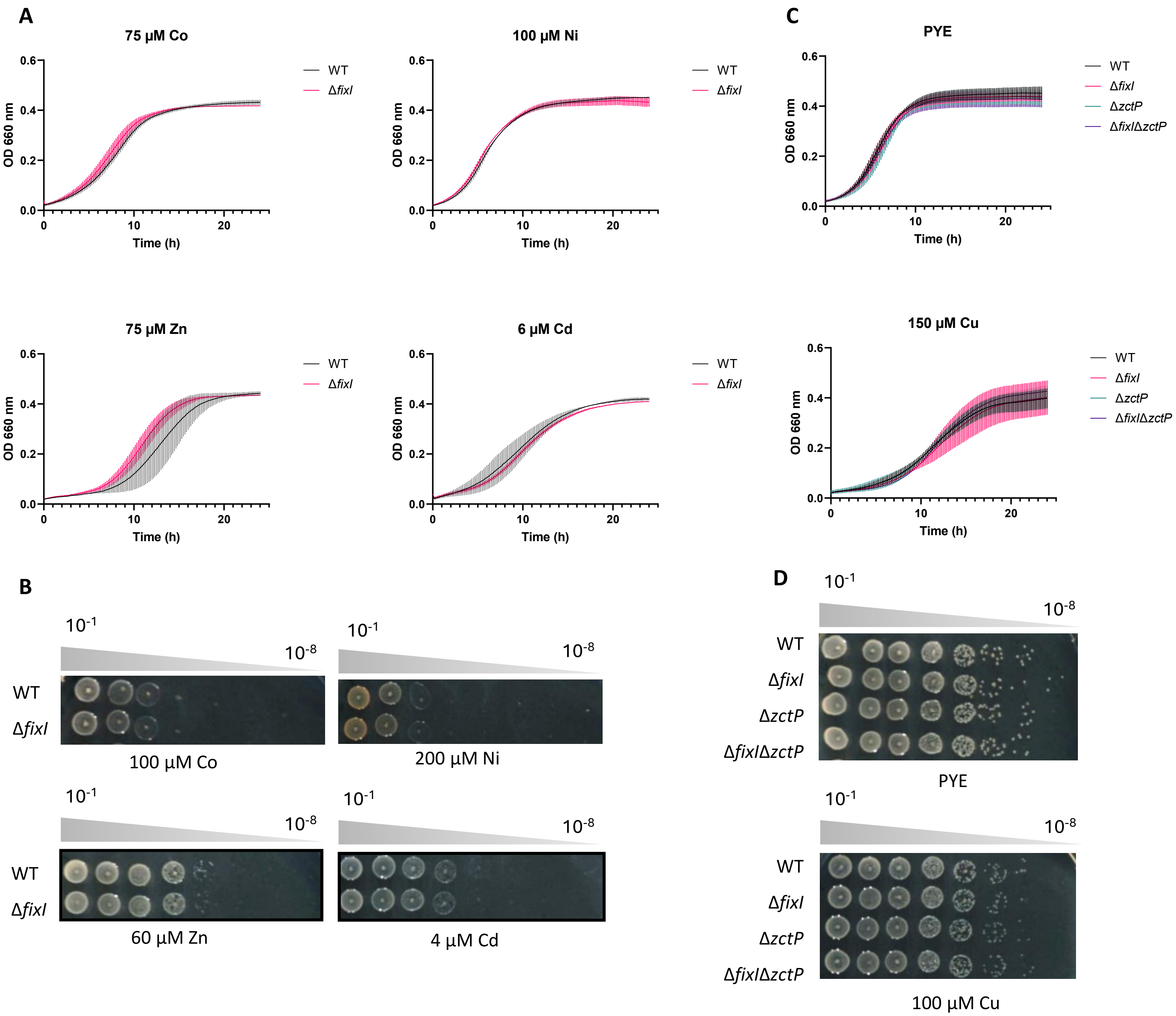


**Figure S2. *C. vibrioides* lacks the P1B-type ATPase required for Cu detoxification. A.** Growth profiles at $OD_{660}$ of WT and Δ*fixI* under Cobalt (Co), Nickel (Ni), Zinc (Zn), and Cadmium (Cd) excess. Mean ± SD, at least three biological replicates. **B**. Viability assay on rich (PYE) under Cobalt (Co), Nickel (Ni), Zinc (Zn), and Cadmium (Cd) excess. Plates were incubated for 48 h at 30 °C before imaging. Biological replicates = 3. **C.** Growth profiles at an absorbance of 660 nm of WT, Δ*fixI*, Δ*zctP,* and Δ*fixI*Δ*zctP* in PYE and moderate Cu stress, Mean ± SD, at least three biological replicates. **D.** Viability assay on rich (PYE) under excess Cu. Plates were incubated for 48 h at 30 °C before imaging. Biological replicates = 3. Whole amino acid sequence alignment of FixI from *C. vibrioides* with its homologs CopA from *Acinetobacter baumannii*, CopA from *Rhodobacter capsulatus*, CopA *Pseudomonas aeruginosa*, CtpA *Mycobacterium tuberculosis,* CopA *Escherichia coli*, CopA *Legionella pneumophila,* and CopA *Salmonella typhimurium*

### Figure S3

**A**

```
CoaT   MRKVVADLNAVEEENMPENHTPDDGHNHAHGGFLNR----ILGGRAEVIFAVLCGICLL---LGWLGPKYGIMSE QFGFGLLLAAYFFGGYFTLREAVE-KISKGQFQIDFLMLVAASGAAILGEWAEGAFLLFLFSVGHALENYAMGRA
ZctP   MAAGIKPLQALHEP-SP-DHGPNTGQ-HSHEGDHARTLGGVLAKNAELVLATGSGAALA---AGWLLSLRA--PG LAPLLCFLLAYGLGGVFTLKEAIE-NLRKRRLAIDSLMLVAAVGAAALGQWAEGALLLFLFSIGHALEHFAMGRA
CtpD   MTALYP----AVEP-AP-AARPARP---RSGGWLWT----VPS----VRWAAAALALFLTGLAAQLLG--A--PQ AVVWTLYLACYVVGGWEPAWVGVR-ALRNRTLDVDLLMIVAAIGAATIGQVFDGALLIVIFATSGALEDVATTRT
PfeT   MNEQVIVQHDPHEP-L------------KTGQKEK----NWAQHGELIAALVSGALIL---AGWLLS--G--HQ VLSITLYLLAFVIGGFAKAKEGIEETLESKTLNVELLMIFAAIGSALIGYWAEGAILIFIFSLSGALETYTMNKS

cons   *          *                 *           .    :  *  .   :     . *    .           *  .:  .**       .:.   :  .   : ::  **:.** *:* :*    :**:*:.:*: . ***   :   ::

CoaT   RNAVAALAGLTPDEALVRRGDKTETVLIENLLVGDIVVVRSNERLPADGFVVKGSSAVNQAPITGESAPVDKLPV DDPEFAAANLDKLTPQTRVFAGSINGSGSLDVQVTKLSGESTLARVVTLVAEAQTRQSPTQNFTKKFEKIFVPCV
ZctP   RRAIEALAKLAPERASVRRGDTVVDVSIEDLEIGDIVLVRPNARIACDGLVIVGQSSVDQAPVTGESVPVDKRPA RDPAL---SFERAGAEHRVFAGTINGAGALDIRVARKASETTLARVVRMVAEAEAQRSPTQQFTDRFERIFVPLV
CtpD   ERSVRGLLDLAPEHATLLGDGSQRVVAAADLRPGDVIVVRPGERISADGTVIGGASEVDQSSITGEPLPAAKDVG D----------------DVFAGTVNGSGALRVEVTREPSQTVVARIVAMVTEASATKATTQLFIEKIEQRYSAGV
PfeT   SRDLTSLMQLEPEEATLMVNGETKRVPVSDLQAGDMIVIKPGERVAADGIIESGSTSIDESALTGESMPVEKNTG D----------------AVFTGTVNRNGSLTVRVTKANEDSLFRKIIKLVESAQSSVSPAQAFIERFENAYVKGV

cons    . : .*  * *:.* :  ..    *   :*  **:::::.. *:..** :  * : ::::.:***. *. *                  **:*::*  *:* :.*::   :: . ::: :* .*.:  :.:* * .::*. :   *

CoaT   IALAFVTSFSF-LILDETAAQSFYRAMAVLVAASPCALAIATPSAVLSGVARAARGGVLIKGGAPLEAMGHLDAI AFDKTGTLTIGEPHLVEITPYGDATETELLQVSAAVEMLSDHPLAQAVVRDVKDRLGDL-PSEASDFANIIGQGV
ZctP   LLAVGLLMLAF-LVIEEPFNVSFYRAMAVLVAASPCALAISVPSAVLSGVARAGRGGVLIKGGGPLENLGALSAM AFDKTGTLTQGKPRLTDVRPEQGVQEHDLLAVAVAVEALSDHPLAGAMVRDGLERLGSRAPGRAEAARSLTGKGV
CtpD   VVATLALLTVP-LMFGADLRSTLLRAMTFMIVASPCAVVLATMPPLLSAIANASRHGVLVKSAVAMERLADTDVV VLDKTGTLTAGEPVISRVTVL--IDGADVLGMAAAAEQFSEHPLGRAIVAAARGRVV-P---EAGDFTALPGRGV
PfeT   LIAVGLLLFVPHFALGWSWSETFYRAMVFMVVASPCALVASIMPAALSLISNGARNGMLVKGSVFLEQLGSVQMI AFDKTGTVTKGQPAVETIRIAEGFSEEEVLEAVYAIETQSSHPLAQAITTYAESRGV-NQSG-YISIEETSGFGV

cons   :  .        : :      :: ***..::.*****:. :  .. ** ::...* *:*:*..  :* :.  . : .:*****:* *:* :  :          ::*    * *  *.***. *:.      *                    * **

CoaT   SAKVDSKVVHIGKTALFESVAGLPLPDDLRGTVEAMSQNGRTTMIVRSGDRYLGAIGLMDTPREDARSVIAALRD LGLKRMMMISGDNQNVANAVAKEVGLDTAFGDLMPEDKVTKIAALKAD-GGVAMVGDGVNDAPAMANATVGIAMG
ZctP   WALVSGKAAIIGKPGLFDGKDAPTMTVSVQALAEDLEAQGRTVVVVSHGERFLGVLGLMDTPRPAAKEVIQRLRR LGLKQIIMLSGDNQTVVSAVGRELGLDEARGGLMPEDKVAIIREMAARNGRVAMVGDGVNDAPAMANATVGVAMG
CtpD   RARVAGHVVEVVSPAAYAGENAA-----VREHCAAIENDGGTAVVVLEDGLPVGVIGLADRLRPDAPAAVMQLAQ LTKHPPMLLTGDNRRAAGRLAEEAGIADVHAELLPDGKAAAVQKLQRDNTHVLVVGDGVNDAPAMAAAHTSIAMG
PfeT   MAEVSGAKWKIGKAGFI-GEEMA--AQFIKQTAPDIIRSGYTIVFVKKDDQIAGCIALKDQIRPEAKEVMEELNR LGIKTA-MLTGDHEDTAQAIANEAGMTTVAAECLPDQKVNEIIRLKEEFGTIAMVGDGINDAPALKAADVGIAMG

cons    * * .   : ...   .          ::     :  .* * :.*   .    * :.* *  *  *  .:  *    *   *  :   :::**:. ..   :..* *:   . .   :*: *.   :   :         : :****:*****:  * ..:***

CoaT   AAGSDVALETADIALMADDLQTLPFAVGLSRKTSRIIRLNLWFSLGVVALLIPATLFG-LGIGPAVLVHEGSTLV VVANALRLLAFKDNRVKSA---
ZctP   AAGSDVALETADVALMADDLSHLPFAVGLSRQTSRIIKQNLWVSLGMVALLIPAALFG-LQIGVAVVFHEGSTLL VVANALRLLAYRDR---TA---
CtpD   RAGADLTVQTADVVTIRDELATVPAVIALARRARRVVIANLVMAGAAITTLVLWDLFGQLPLPLGVAGHEGSTIL VALNGLRLLSNRAWISPGATPT
PfeT   -GGTDVALETADMVLMQNDLKKLVNMCRLSRKMNRIIKQNIVFSLAVICLLICANFLQVMELPFGVIGHEGSTIL VILNGLRLLK------------

cons    .*:*::::***:. : ::*  :      *:*:   *::   *: .: . :   *:    ::  : :   .*   *****:: *   *.****
```

**B**

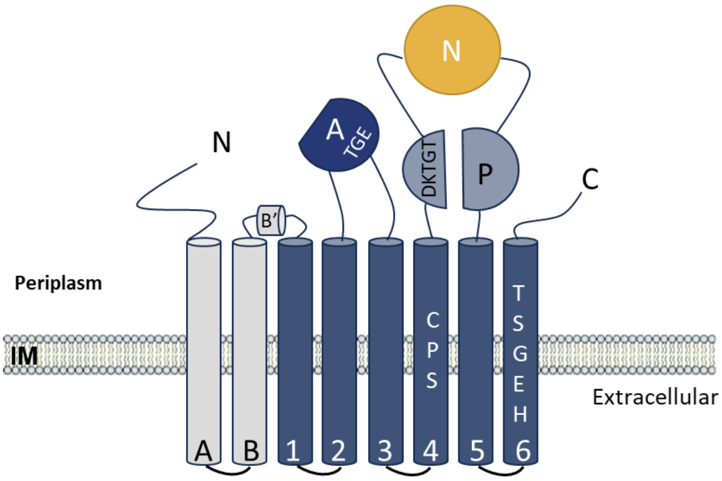


**Figure S3. ZctP seems to belong to the P1B4 subclass of P-type ATPases. A.** Whole amino acid sequence alignment of ZctP *C. vibrioides* with its homologs CoaT *Sulfitobacter sp*. NAS14-1, CtpD *Mycobacterium smegmatis* and PfeT *Bacillus subtilis*. **B.** Schematic topology of P-type ATPases showing features unique to PIB4-ATPase of ZctP. Key residues for PIB4-ATPases are highlighted.

### Figure S4

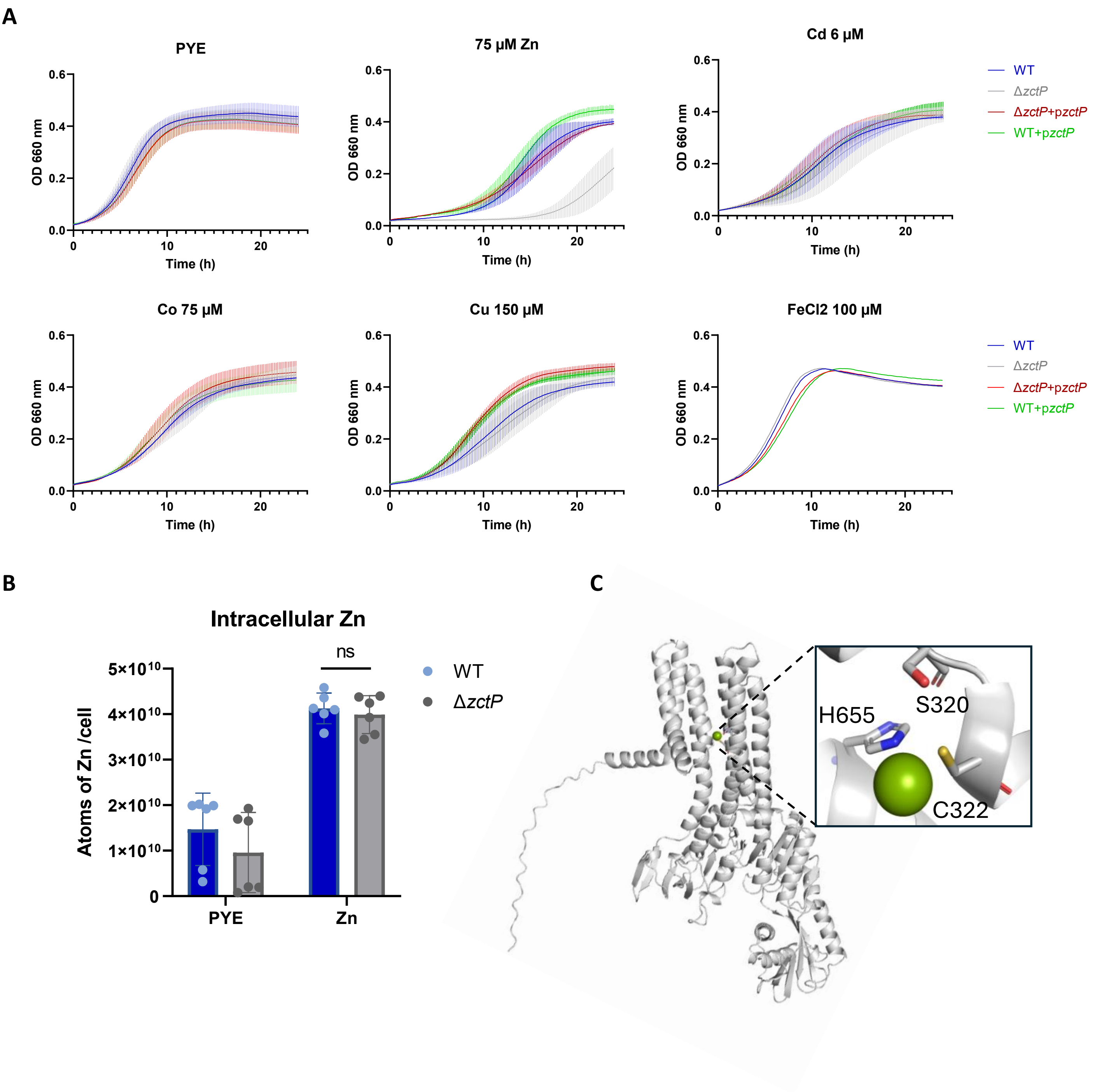


**Figure S4. ZctP is specifically required for Zn homeostasis. A.** Growth profiles at $OD_{660}$ of WT, Δ*zctP, complemented strain (*Δ*zctP*+ p*zctP) and overexpressed strain (*WT+ p*zctP)* under excess Cobalt (Co), Nickel (Ni), Zinc (Zn), and Cadmium (Cd), Cobalt (Co), Cu (Cu) and Iron (Fe) excess. Mean ± SD, at least three biological replicates except the replicate with excess Fe. **B.** Prediction by Alphafold of ZctP 3D structure with a potential Zn binding site. The model was visualized using Pymol. **C**. The Number of Cu atoms per cell of the WT and ΔzctP strains in the control condition (PYE) was exposed to $ZnSO_4$ excess for 30 min. Mean ± SD, at least three biological replicates. P-values were calculated using a t-test (*$p < 0.05$, ** $p < 0.01$, *** $p < 0.001$, and ****$p < 0.0001$) **(Table S4)**.

### Figure S5

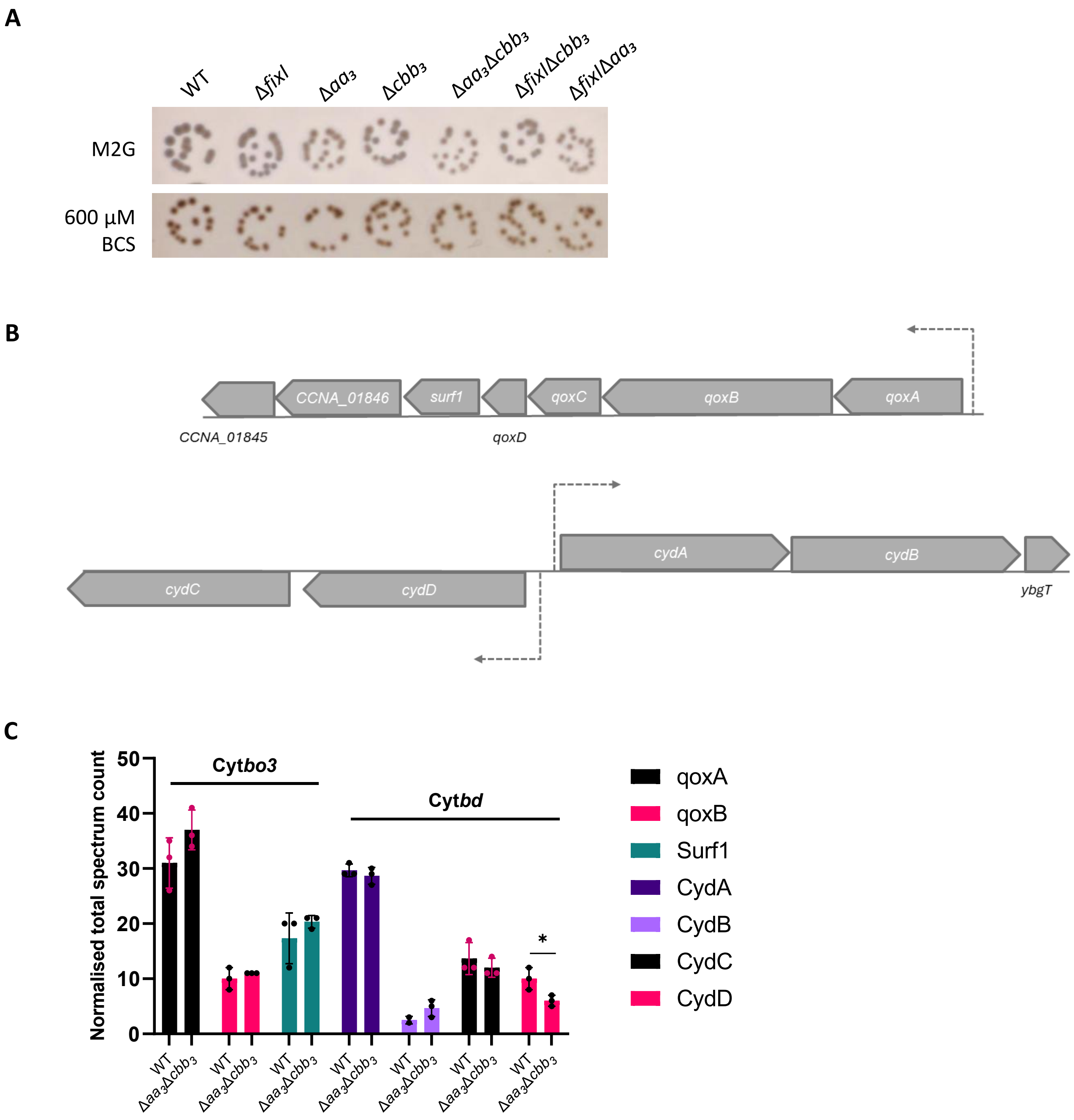


**Figure S5. *C. vibrioides* possesses two quinol oxidases: cytochrome *bd*-type quinol oxidase (Cyt*bd*) and cytochrome *bo3* quinol oxidase (Cyt*bo3*). A.** Growth and NADI phenotypes of colonies of *C. vibrioides* wild-type (WT) and mutant strains. Cells were grown aerobically at 30°C on a minimal medium (M2G) and M2G supplemented with 600 µM BCS (Cu chelator), and the presence of Cox activity was visualized by NADI staining (see Materials and Methods). **B.** Genomic organization of the Cyt*bo3* and Cyt*bd* operon, respectively. **C.** Normalized total spectrum count of peptides from Cyt*bo3* and Cyt*bd* subunits in the WT and $\Delta aa_3\Delta cbb_3$ strains grown in PYE medium, as measured by LC-MS. Individual values and means are shown. **C.** Normalized total spectrum count of peptides from Cyt*bo3* and Cyt*bd* subunits in the WT and $\Delta aa_3\Delta cbb_3$ strains grown in PYE medium, as measured by LC-MS. Individual values and means are shown. P-values were calculated using a t-test (*$p < 0.05$, ** $p < 0.01$, *** $p < 0.001$, and ****$p < 0.0001$) **(Table S4)**.

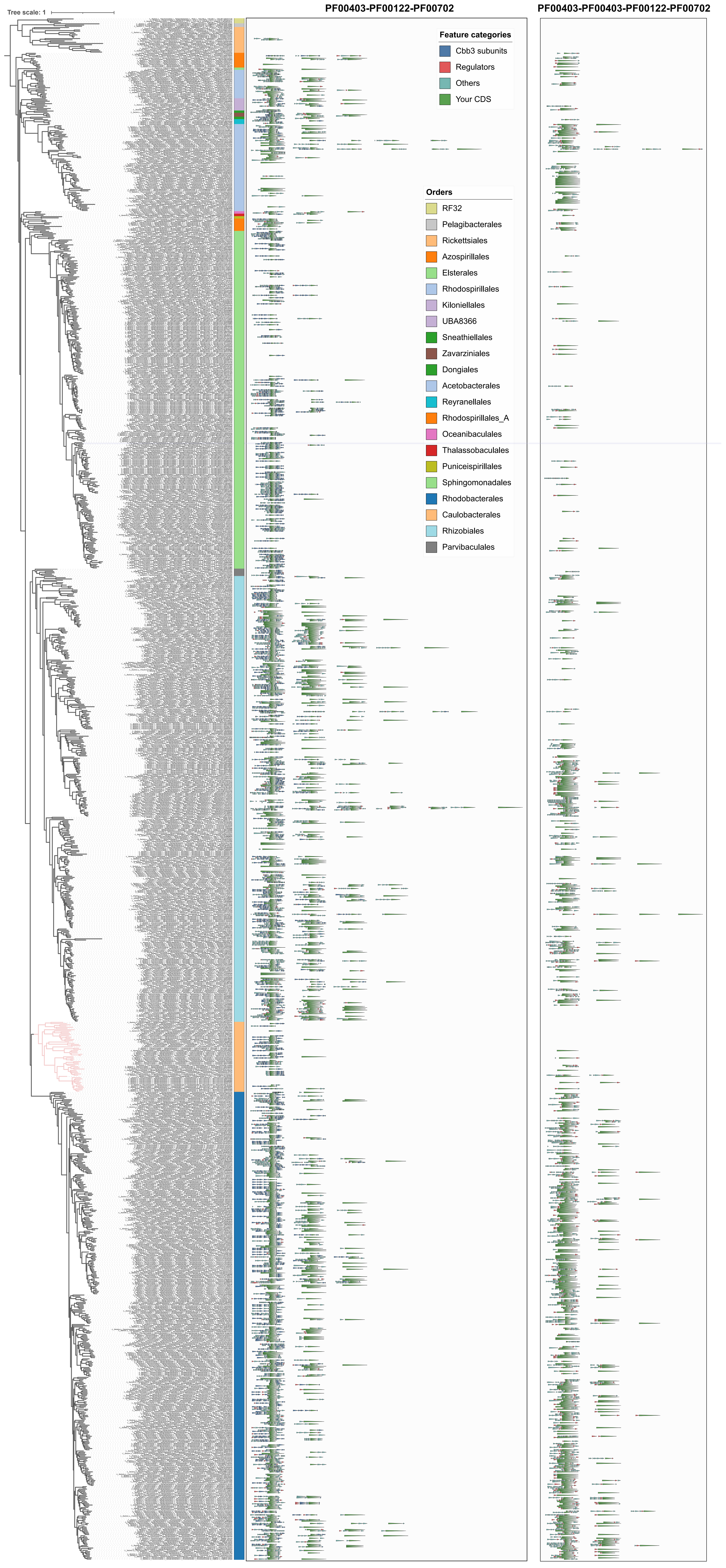
Tree scale: 1
PF00403-PF00122-PF00702
PF00403-PF00403-PF00122-PF00702
Feature categories
Cbb3 subunits
Regulators
Others
Your CDS
Orders
RF32
Pelagibacterales
Rickettsiales
Azospirillales
Elsterales
Rhodospirillales
Kiloniellales
UBA8366
Sneathiellales
Zavarziniales
Dongiales
Acetobacterales
Reyranellales
Rhodospirillales_A
Oceanibaculales
Thalassobaculales
Puniceispirillales
Sphingomonadales
Rhodobacterales
Caulobacterales
Rhizobiales
Parvibaculales

### Figure S6

**Figure S6. Genomic context of P-type ATPases across Alphaproteobacteria**. GTDB phylogenetic tree of 1,255 Alphaproteobacterial genomes, with taxonomic orders color-coded as indicated. To the right, the genomic context is shown for all proteins containing the two domain architectures of interest: PF00403–PF00122–PF00702 (left panel) and PF00403–PF00403–PF00122–PF00702 (right panel). Genes are colored by predicted function: $cbb_3$-type cytochrome c oxidase subunits (blue), regulatory elements (red), other genes (gray), and the ATPase of interest (green)

### Figure S7

**A**

```
    PccA C. vibrioides  MKTLTL-LGVCAAALALTAAAPAPKV------TAADAWCRPAPTGAMAGGCYVTLT-A-GVDDRLVAVETTAAARGEIHT
     PccA R. capsulatus  MIKGFA-APLCALMLALMPVSGHAHEAQAGALTIIHPSIPAVMPGAQAAAGYFTIVNAGDSADRLTAVEVDFAAMAMLHK
PCuAC R. sphaeroides  MTPFQAILAAAAVTFALPAFAD-SEL------AVTDAYARVASPAAQSGAIFMVIENHGASDDRLVSAATDAAARVELHT
                      *      .* :** . :  ..     :  ..   . ..* :.. :..:     ***.:. .  **  :*.

    PccA C. vibrioides  MSM-DGGVMRMRKLADGLALPAGKAVALKPGADHIMVIGPKIALTEGAQVPLTLKFQKAKPVKVTAVVKMPPK-P---AM
     PccA R. capsulatus  TETDAAGVARMLDV-DAVEVPAGATVTFAPGGLHVMFMGLSAPIAEGVMLPGTLVFEHAGKVPVEFMVTTAGKAP---DM
PCuAC R. sphaeroides  HLAGQDGVMQMVEVKEGFPVPAHGSHALARGGDHVMLMGLTKPLKEGDEIALTLTFESGKVLEVAAPVDTSREAPMGDHM
                           ** :* .: :.. :**  : ::  *. *:*.:* . .: **  :. ** *: .  : *  *  . : *   *

    PccA C. vibrioides  SGMH--H
     PccA R. capsulatus  SGHAMP-
PCuAC R. sphaeroides  KGMTHGN
                      .*
```

**B**

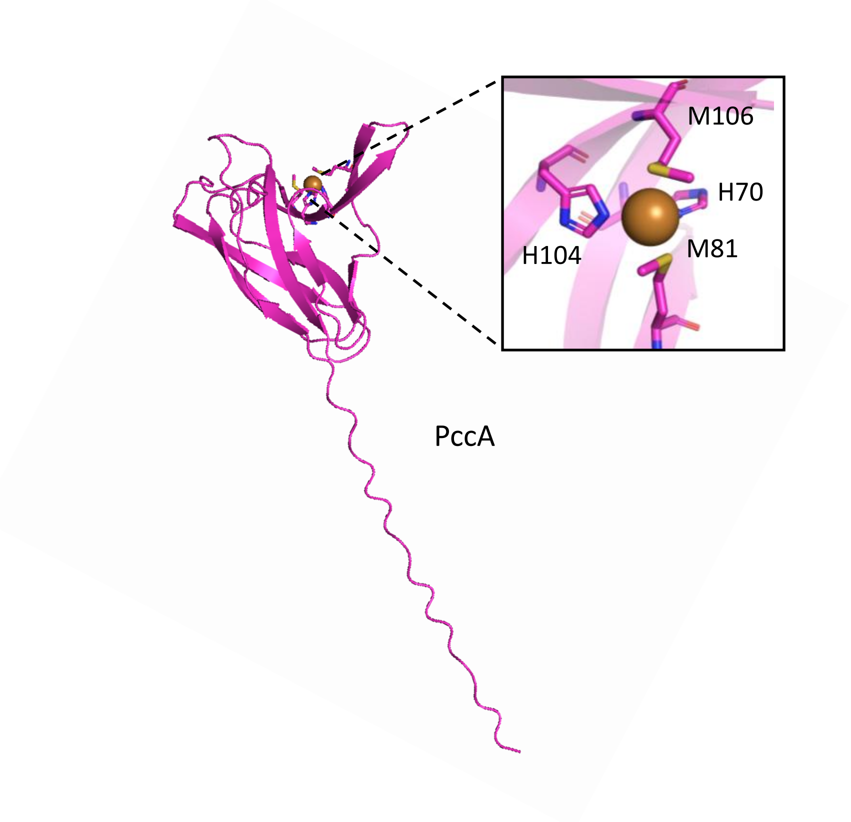


**Figure S7. PccA from *C. vibrioides* belongs to the PCuAC family. A.** The conserved H(M)$X_{10}$M$X_{21}$HXM motif from PccA sequence alignments from *C. vibrioides*, *R. capsulatus,* and *R. sphaeroides*. **B.** Alphafold predicted structure of *C. vibrioides* PccA. PccA exhibits a cupredoxin-like fold, a structural motif common in Cu-binding proteins. This fold involves two histidine and two methionine residues that could coordinate a Cu(I) ion (His 70, Met 81, His 104, and Met 106).

### Figure S8

**A**

```
Sco1 C. vibrioides   ----------------------------MPRHRLVLILACVVGLAV----------------------------AVGLAW
Sco2 C. vibrioides   ------------------------------------------MRAIALTLALLAATPLAGCGEKAAQDAAGAVIKIG---
SenC R. capsulatus   MTKLYAGVAAAAIAALLAGSAAWVFLGRSEERFAGCGANQVAGGA-----------------------------------
PrrC R. sphaeroides  -------------------------------------MNVSSKTA------ALAATAAVVVVVGISAAVTLVPHETDRFA
                                                                 *

Sco1 C. vibrioides   NVGVFRSEPAVTVGGPFELVDQNGAPTSEKALKGKWSAVFFGFTYCPDVCPGTLQGLAAATDQLGPKAKDFQIVFISIDP
Sco2 C. vibrioides   --------------GPFQLTDMNGKPVTEKSLLGKPTAVFFGFTYCPEVCPTTLTEMTAWLKALGKDADKLNVVLITVDP
SenC R. capsulatus   ------------IGGPFTLVDQEGRTVTDREVLAKPSLVYFGYTFCPDVCPFDMARNAQAVDILTEWGIEVTPVFISIDP
PrrC R. sphaeroides  ACRKGTGSASAQIGGPFTLISETGATVTDRDVITKPSLVYFGYSYCPDVCPIDSTRNAAAVDLLAERGHDVTPVFISVDA
                                   *** *  .  *  ..::: :  *  : *:**:::**:***      :   . *   . ..  *:*::*.

Sco1 C. vibrioides   ARDTVKQMKAYLSA--PYVPKATIGLTGTQAQVDAAAKAYRVYHAKV--GDGVDY-TMDHSTAIYLMDPKGRFKTVIPYN
Sco2 C. vibrioides   ERDTPAQLKEYLSNFDPRIQ----GFTGTPDAIAKTARAYRVYYQKVPLDGG-GY-TIDHSSAIYLFDAKGRF--VSPIA
SenC R. capsulatus   KRDTPEQLKFFAEAIHPDTI----ALTGTEAQVKAASQAYKTFY-RVQESDD-DYYLIDHSTFTYFMLPGTGF--VDFF-
PrrC R. sphaeroides  ARDTPPVLTEFTDLMSPKMI----GLTGTPEQIDAAVKAYRAYYLIRNPGDP-AT-LVDHSTQTYLMDPKLGF--LDFYD
                      ***   :. : .   *       .:***   :  : :**:.::     ..      :***:  *:: .   *  :

Sco1 C. cvibrioides  L--P-------------PDE--IA--RR-IKDAVREG----------
Sco2 C. vibrioides   YQ-A-------------PQD--RA-LGQ-LRDLLK------------
SenC R. capsulatus   KREDTPEQIAERISCFANDSHVSTSFDARAQKSYQASRGKQMGDNHG
PrrC R. sphaeroides  RDA--TPEMVADSVGCFLDA---LQTPG-DTPAA-----GNGN----
                                       :
```

**B**

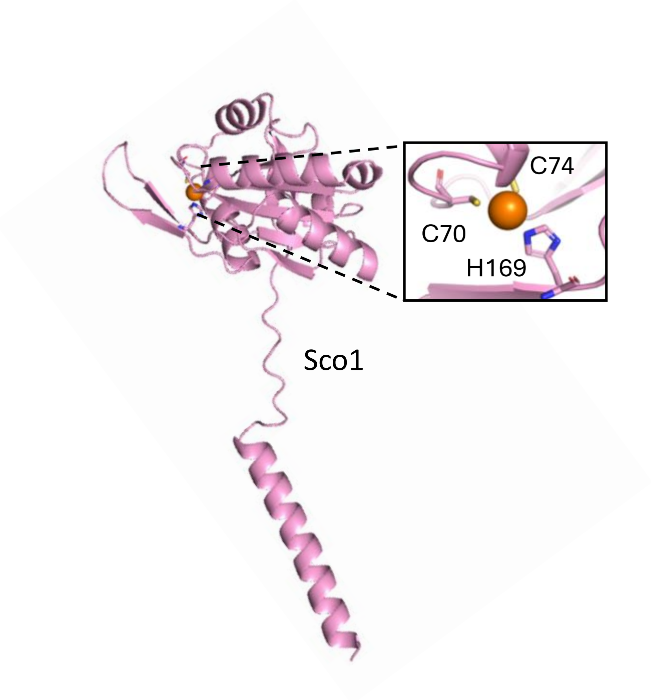


**C**

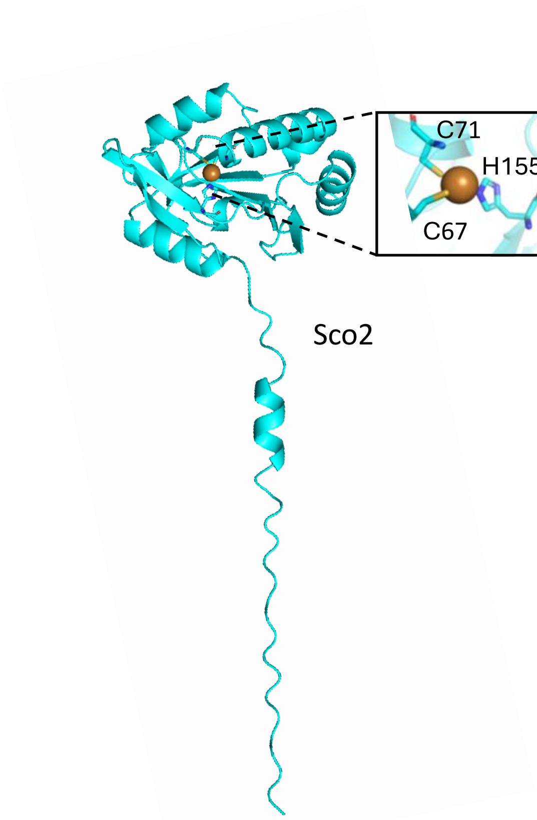


**Figure S8. *C. vibrioides* possesses two paralogs of the Sco family Cu chaperone. A.** Amino acid sequence alignment of Sco1 and Sco2 from *C. vibrioides* with its homologs SenC from *R. capsulatus* and PrrC from *R. sphaeroides*. The canonical CXXXCP motif, which is proposed to bind Cu, is highlighted in light orange in addition to the conserved H residue. **B, C.** Alphafold prediction of Sco1 and Sco2 3D structure, respectively, highlighting the conserved thioredoxin-like fold. The conserved key residues possibly involved in Cu binding are C70, C74, and H169 for Sco1 and Cys 67, Cys 71, and His 155 for Sco2

### Figure S9

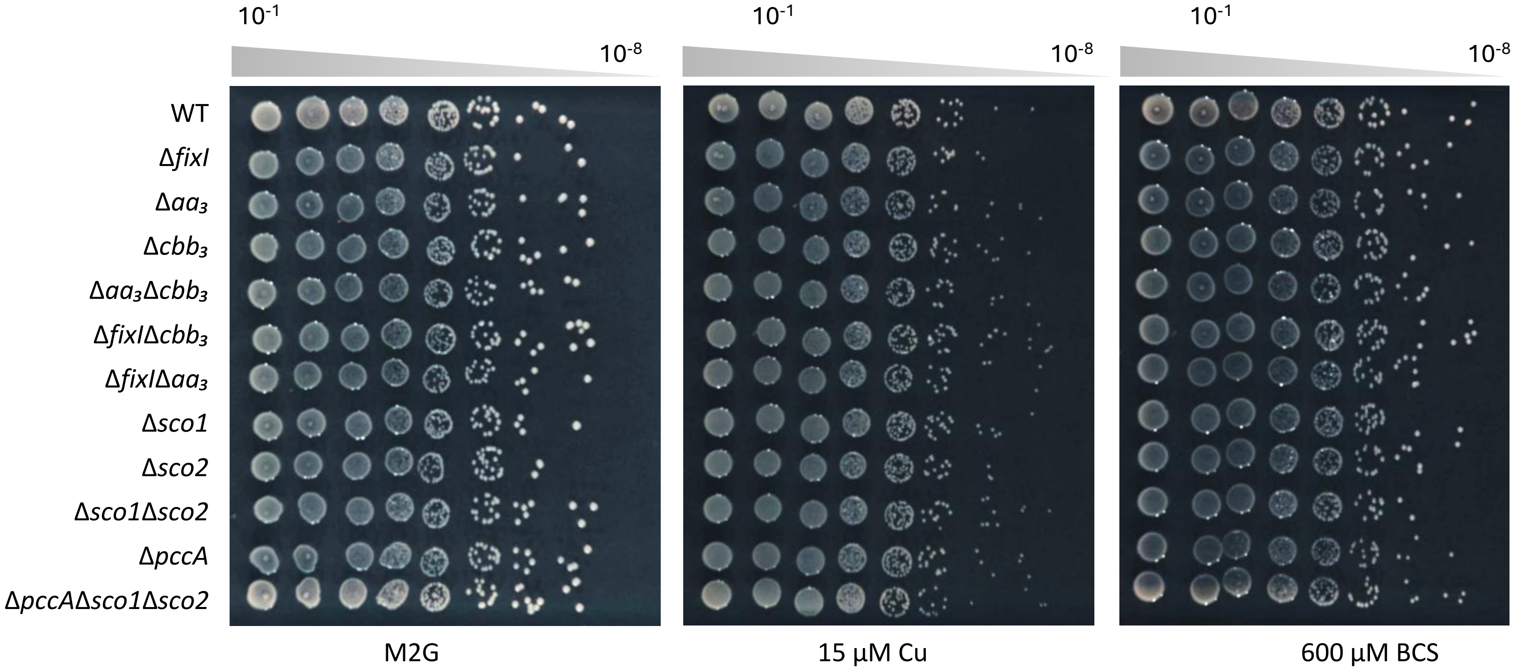


**Figure S9. The potential PccA, Sco1, and Sco2 Cu chaperones are not required for Cu resistance in C. vibrioides.** Viability assays were performed on minimal medium (M2G) under control conditions, Cu-excess (15 µM $CuSO_4$), and Cu-deficiency (600 µM BCS). Plates were incubated for 48 h at 30 °C before imaging. Biological replicates (n=3).

### Figure S10

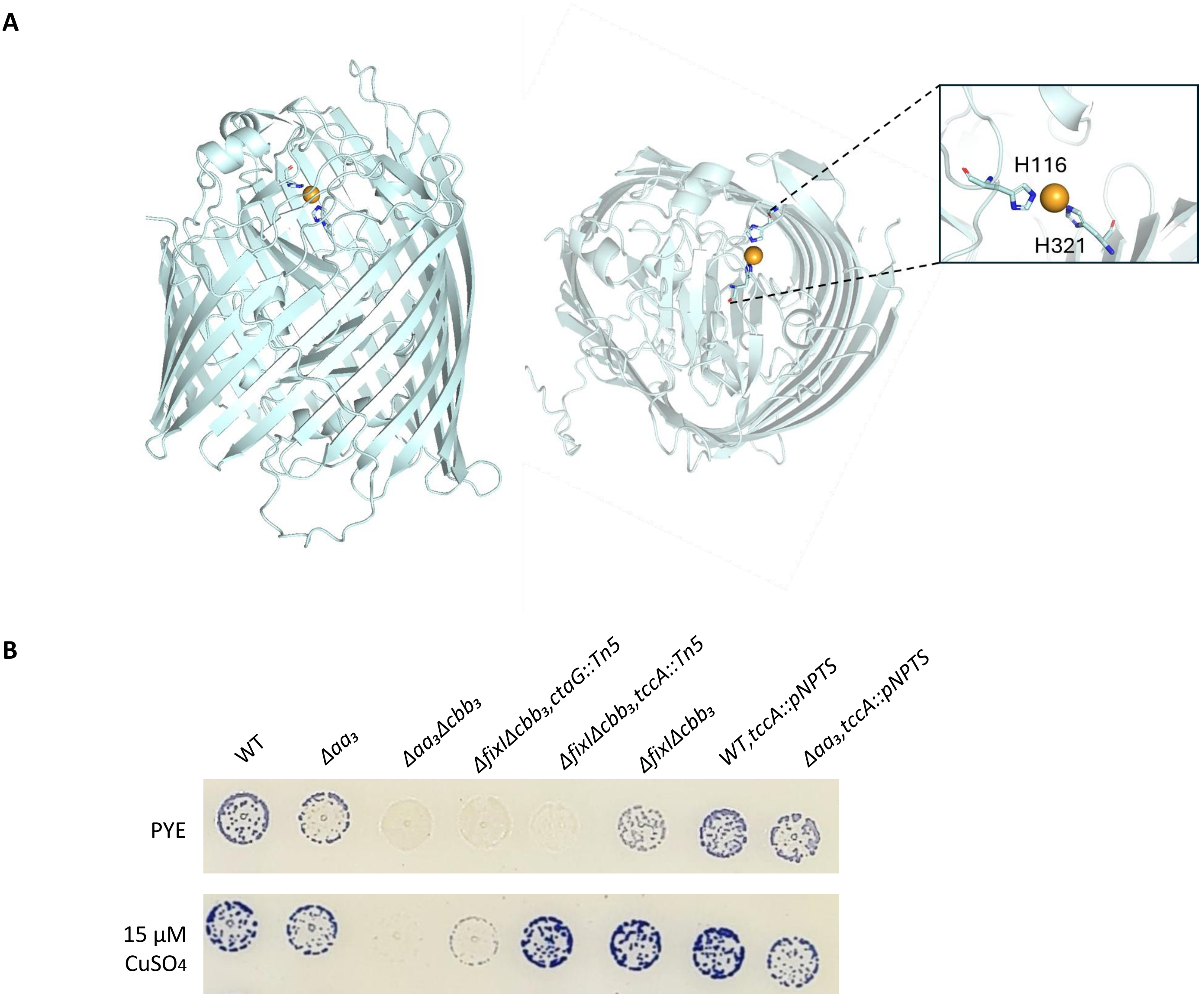


**Figure S10. TccA has a potential Cu binding site and is specifically required for *aa*$_3$-Cox. A.** Structure prediction of TccA by AlphaFold with a proposed Cu binding site composed of two His residues (H116 and H321. **B.** Growth and NADI phenotypes of colonies of *C. vibrioides* WT and mutant strains. Cells were grown aerobically at 30°C on a PYE medium and PYE supplemented with 15 µM $CuSO_4$ , and the presence of Cox activity was visualized by NADI staining. Colonies that contain WT levels of Cox activity turn dark blue within a few seconds (NADI$^+$), while those that have low or no Cox activity show lighter blue (NADI$^{slow}$) or no blue staining (NADI$^-$) upon longer exposure, respectively. (entire plate in Fig. S12B)

### Figure S11

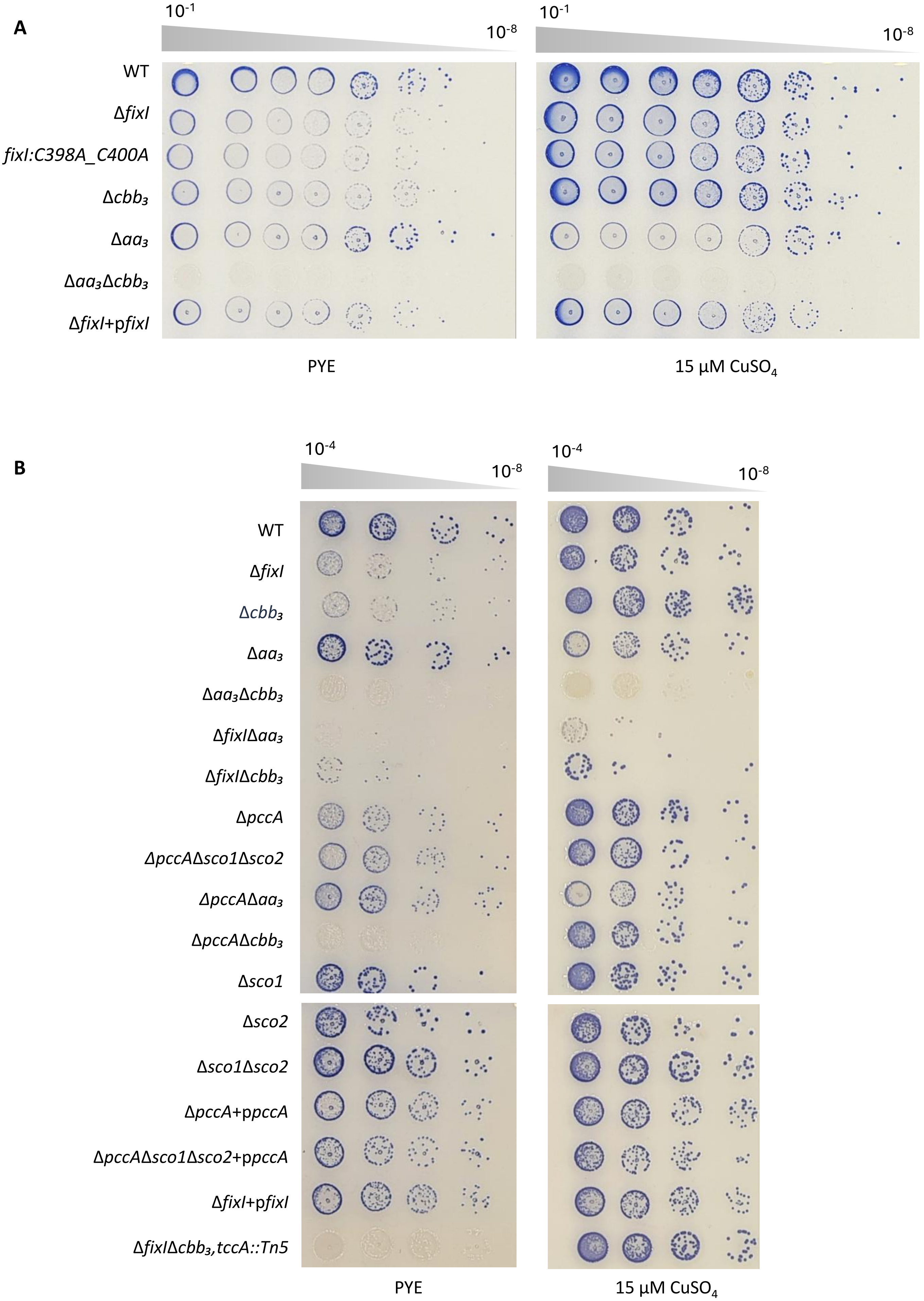


**Figure S11. Cox activity of the different *C. vibrioides* strains used in this study visualized by NADI stain in control rich medium and in the presence of 15 μM $CuSO_4$**

### Figure S12

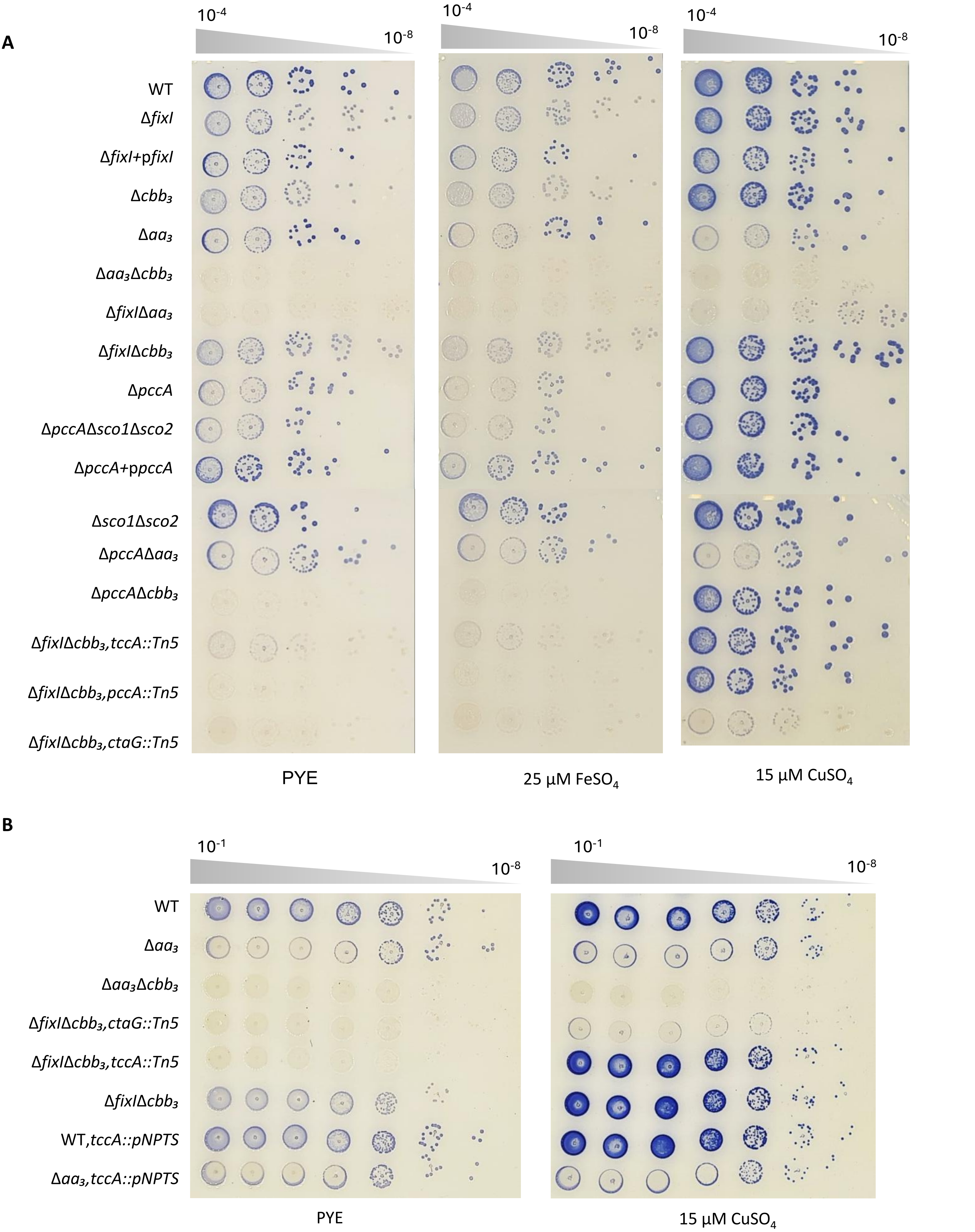


**Figure S12. Cox activity of the different *C. vibrioides* strains used in this study visualized by NADI stain in control rich medium and in the presence of 15 µM $CuSO_4$ and/or 25 µM $FeSO_4$.**
