## Supporting text for "Unraveling the pathway of Copper Delivery to Cytochrome *c* oxidases in the Free-Living Bacterium *Caulobacter vibrioides*"

### Additional information 1

**The P1B4-type ATPase ZctP seems to be required for Zn homeostasis in *C. vibrioides***

To classify ZctP within the P-type ATPase family, an amino acid alignment was performed using T-COFFEE, comparing ZctP from *C. vibrioides* with other characterized P1B4-type ATPases, CoaT from *Sulfitobacter sp. NAS14-1,* a Co-specific transporter(41), CtpD from *Mycobacterium smegmatis*, mediates Co and Ni efflux (42) and PfeT from *Bacillus subtilis* functions as an Fe(II) exporter (43). P1B4-type ATPases are characterized by a distinctive SPC metal-binding motif in the transmembrane region 4 and a HEG(G/S)T metal-binding motif in the transmembrane region 6 (Fig. S3A). This analysis suggests that ZctP belongs to the P1B4 subclass of P-type ATPases. The topology and TM-helix positions were predicted using TOPCONS (44). ZnctP contains eight transmembrane helices: MA and MB, along with a shorter segment MB', followed by helices M1 to M6 (Fig. 1B). MA, MB, and MB' are membrane-associated regions at the N-terminus that may help stabilize the protein or assist in metal binding. Structural analysis shows that the P1B4-type ATPase architecture resembles that of other P-type ATPases, featuring three cytosolic domains: A (actuator), N (nucleotide-binding), and P (phosphorylation), which includes the conserved DKTGT motif where the aspartate residue is phosphorylated during the transport cycle (Fig. S3B) (45, 46).

To investigate the role of ZctP in *C. vibrioides* physiology, the growth profile of the clean *zctP* knockout strain (Δ*zctP*) was monitored in a control-rich medium (PYE) and under various metal stress conditions. Both the wild-type (WT) strain and Δ*zctP* displayed similar growth in PYE, indicating that ZctP is not essential for the survival of *C. vibrioides* under these conditions. When exposed to moderate Zn stress, the growth of the Δ*zctP* mutant was impaired (Fig. S4A). This increased sensitivity was reversed by introducing an exogenous copy of *zctP* under the constitutive plac promoter (Δ*zctP*+p*zctP)*, confirming that the observed phenotype was not due to a polar effect (Fig. S4A). The WT strain carrying an exogenous copy of *zctP* under the constitutive plac promoter (WT+p*zctP*), showed a growth profile similar to that of the WT and complemented strains, suggesting that ZctP is necessary for Zn homeostasis under the tested conditions (Fig. S4A). No sensitivity was observed when the mutant strain was exposed to Cd, Co, Cu, or Fe (Fig. S4A).

To determine whether the observed sensitivity of the Δ*zctP* strain under excess Zn was due to the accumulation of toxic Zn, the intracellular Zn concentrations of the WT and Δ*zctP* strains were measured using an atomic absorption spectrometer (AAS). No significant difference in intracellular Zn levels was found between the WT and Δ*zctP* strains, suggesting that the sensitivity is not caused by Zn accumulation (Fig. S4B). AlphaFold 3 prediction of the ZctP Zn-binding site also identified residues S321, C323, and H655 as key components of the binding site (Fig. S4C).

Altogether, *C. vibrioides* possesses a P1B4-type ATPase that appears to play a role in maintaining Zn homeostasis. While current evidence points to Zn as the likely substrate of ZctP, its exact function, transport directionality, and role in maintaining metal homeostasis remain to be elucidated in further biochemical and physiological studies. The study by Marques *et al.* (47) identified and characterized two Resistance-Nodulation-Division (RND) efflux systems in *C. vibrioides*: CzrCBA and NczCBA that mediate resistance to heavy metals. The *czrCBA* operon is strongly induced by Cd and Zn, and to a lesser extent by Co and Ni, while *nczCBA* is primarily responsive to Ni and Co. Functional studies showed that deletion of *czrA* increased sensitivity to Cd, Zn, and Co, but not Ni, suggesting a specific role in Zn and Cd detoxification (47). Notably, the *zctP* gene is located in the same operon as *czrCBA*, further suggesting a coordinated function in Zn resistance and export.

### Additional information 2

**PccA, Sco1 and Sco2 are not required for Cu resistance in *C. vibrioides***

Previous studies have shown that purified PccA from *C. vibrioides* binds Cu *in vitro*. Additionally, in this study, *in silico* analysis and AlphaFold predictions suggested that both Sco1 and Sco2 have the potential to bind Cu. To determine whether these Cu chaperones are essential for Cu homeostasis, the growth of the *ΔpccA, Δsco1, Δsco2, Δsco1*Δ*sco2,* and *ΔpccA*Δ*sco1*Δ*sco2* mutants was assessed under Cu-depleted and Cu-excess conditions. CFU was determined on minimal growth medium (M2G) supplemented with moderate Cu stress (15 μM) to evaluate strain viability. Serial dilutions were prepared, and each dilution was spotted onto M2G plates containing excess Cu. None of the tested mutants (*ΔpccA, Δsco1, Δsco2, Δsco1*Δ*sco2,* and *ΔpccA*Δ*sco1*Δ*sco2*) exhibited sensitivity to Cu excess (Fig. S9). Similarly, no growth defects were observed when Cu was chelated using BCS. These findings indicate that none of these chaperones is crucial for Cu homeostasis in *C. vibrioides*. The viability of strains lacking one or both Cox enzymes was assessed under the same conditions, and their growth profiles were comparable to the WT strain profile.
